# Regulation of late-acting operons by three transcription factors and a CRISPR-Cas component during *Myxococcus xanthus* development

**DOI:** 10.1101/2023.07.26.550624

**Authors:** Shreya Saha, Lee Kroos

## Abstract

Upon starvation rod-shaped *Myxococcus xanthus* bacteria form mounds and then differentiate into round stress-resistant spores. Little is known about the regulation of late-acting operons important for spore formation. C-signaling has been proposed to activate FruA, which binds DNA cooperatively with MrpC to increase transcription of many genes. We report that this model can explain regulation of the *fadIJ* operon involved in spore metabolism, but not that of the spore coat biogenesis operons *exoA-I*, *exoL-P*, and *nfsA-H*. Rather, a mutation in *fruA* increased the transcript levels from these operons early in development, suggesting negative regulation by FruA initially, and a mutation in *mrpC* affected transcript levels from each operon differently. FruA bound to all four promoter regions *in vitro*, but strikingly each promoter region was unique in terms of whether or not MrpC and the DNA-binding domain of Nla6 bound, and in terms of cooperative binding. Furthermore, the DevI component of a CRISPR-Cas system is a negative regulator of all four operons, based on transcript measurements. Our results demonstrate complex regulation of sporulation genes by three transcription factors and a CRISPR-Cas component, which we propose thwarts viral intrusion while making spores suited to withstand starvation and environmental insults.

## Introduction

The Gram-negative soil bacterium *Myxococcus xanthus* provides an attractive model system to study signal-induced gene regulation and bacterial community behavior (1). Upon nutrient depletion, cells move on solid surfaces and form mounds, within which some of the rod-shaped cells differentiate into round, stress-resistant spores. During this multicellular developmental process of fruiting body formation, a majority of the population undergoes lysis, while some cells remain outside of fruiting bodies as peripheral rods (2, 3). Transcriptomic analyses indicate that approximately 1500 (4, 5) to 4000 (6) genes (i.e., 20-50% of the genes) are up- or down-regulated during *M. xanthus* development.

A signal-responsive gene regulatory network (GRN) governs the developmental process of *M. xanthus* (7). Starvation triggers production of the intracellular secondary messenger molecules (p)ppGpp (8, 9) and c-di-GMP (10), which lead to production of the extracellular A- and C-signals (9, 11–14) and exopolysaccharide (10), respectively. Other signals are implicated in the developmental process, but are less well-characterized in terms of their identity and/or their impact on the GRN (15).

Among the better-characterized developmental signals of *M. xanthus*, early studies showed that C-signal acts later than (p)ppGpp and A-signal (9, 11, 16, 17). The exchange of C-signal between cells required their movement into alignment, possibly into contact (18–20). An increasing level of C-signaling was proposed to ensure that cells first build mounds and then form spores, resulting in multicellular fruiting bodies (21–23). The C-signal appears to be a proteolytic fragment of the CsgA protein (24–26) and/or diacylglycerols produced by cardiolipin phospholipase activity of full-length CsgA (27), but the mechanism(s) of signal transduction remain to be elucidated. C-signaling appears to posttranslationally activate the transcription factor FruA (28, 29) (Fig. 1), but the mechanism is unknown. FruA is similar to response regulators of two-component signal-transduction systems (30). Response regulators are typically activated by phosphorylation by a histidine kinase (28). However, for several reasons, phosphorylation is unlikely to be the mechanism by which FruA is activated in response to C-signaling (29, 31).

**Figure 1.**
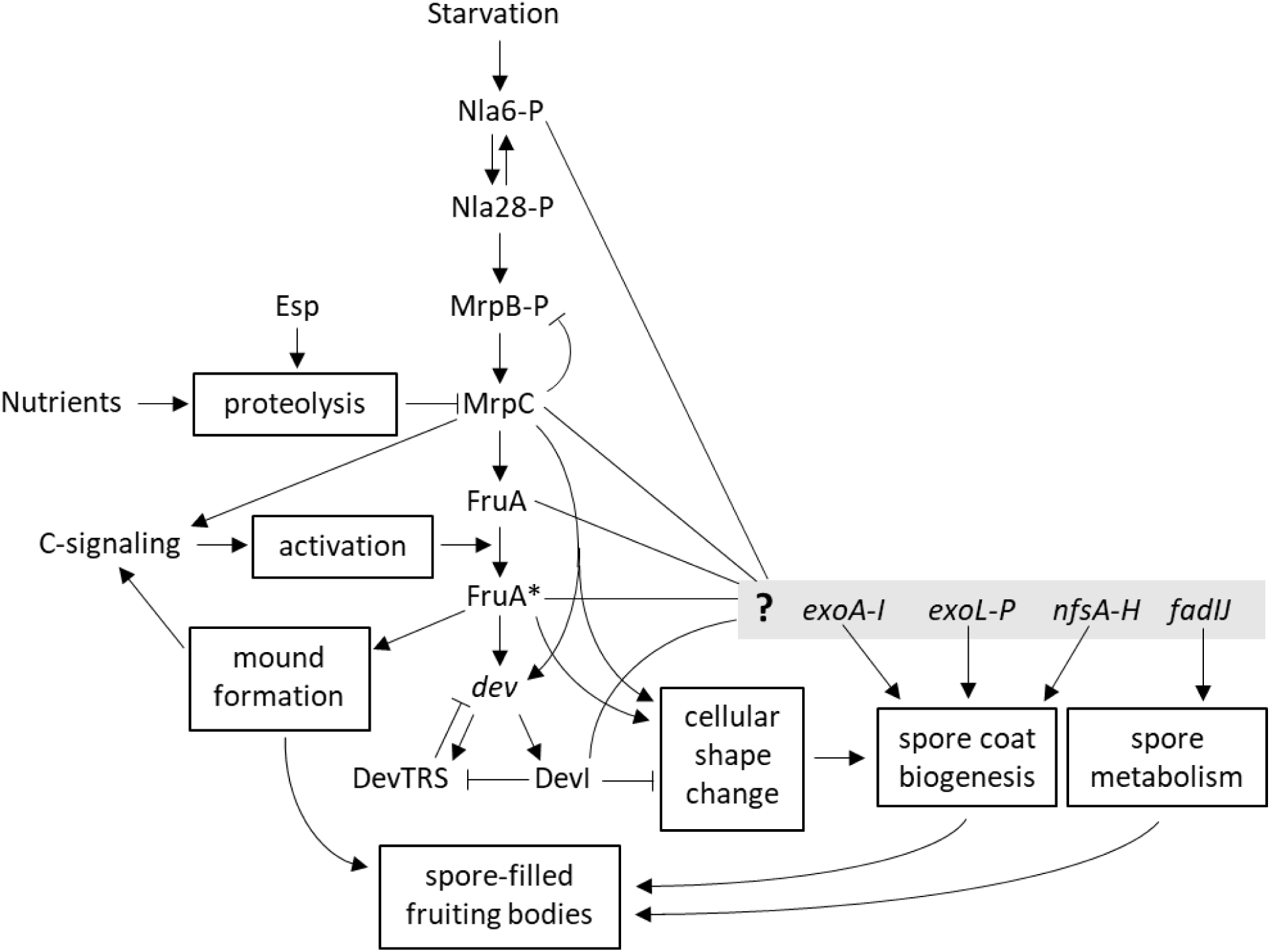
Model of the gene regulatory network governing formation of *M. xanthus* fruiting bodies. Starvation increases the level of phosphorylated Nla6 (Nla6-P), Nla28-P, MrpB-P, and MrpC in a cascade of transcription factors (7, 32, 35). Nutrients (38, 39) and Esp signaling (36, 37) can negatively regulate MrpC *via* proteolysis, and MrpC negatively autoregulates by competing with MrpB-P for binding to the *mrpC* promoter region (35). MrpC positively regulates C-signaling (97) and *fruA* transcription (33), leading to production of unactivated FruA. C-signaling activates FruA to FruA* by an unknown mechanism (28, 29). FruA* positively regulates mound formation (30), which enhances C-signaling (18–20), forming a positive feedback loop. FruA* also positively regulates the *dev* operon (98) together with MrpC (42), thus forming a feed-forward loop that has been speculated to also increase transcription of genes involved in cellular shape change (29). DevTRS proteins negatively autoregulate, decreasing DevI, which negatively regulates cellular shape change (52, 53) and may inhibit negative autoregulation by DevTRS (29). Here, we investigate possible regulation by Nla6-P, MrpC, FruA or FruA*, and DevI (lines to gray box with **?**) of late-acting operons involved in spore metabolism and spore coat biogenesis.

Transcription of the *fruA* gene is regulated by a cascade of starvation-responsive transcription factors that includes MrpC (7, 32) (Fig. 1), which binds upstream of the *fruA* promoter and stimulates transcription (33, 34). In addition to transcriptional control of the *mrpC* gene by starvation (35), the level of MrpC protein is regulated by proteolysis (Fig. 1), slowing the pace of development under the normal conditions of starvation (36, 37) and halting development if sufficient nutrients are added back before rods become committed to spore formation (38, 39). An N-terminally truncated form of MrpC called “MrpC2” was proposed to be important for development (34), but recent work indicated that MrpC2 is likely an artifact of the method of cell lysate preparation and that full-length MrpC is the active form *in vivo* (35), at least in part due to a threonine/serine motif near its N-terminus (40). Both MrpC and MrpC2 bind to DNA *in vitro* (34, 35, 41).

*In vitro*, MrpC2 and FruA bind cooperatively to the promoter regions of many developmentally-regulated genes (41). Mutational analyses of five such promoter regions (*dev*, *fmgA*, *fmgBC*, *fmgD*, and *fmgE*) suggest that cooperative binding of MrpC and FruA to a site just upstream of the promoter stimulates transcription *in vivo*, forming a feed-forward loop (Fig. 1, only *dev* is shown), whereas binding to nearby sites can increase or decrease transcription (31, 42–45). The different arrangements of binding sites with different affinities for MrpC and FruA may explain the observed differences between genes in terms of their spatiotemporal patterns of expression and their extent of dependence on C-signaling for expression (7, 17, 46). The levels of MrpC, FruA, and CsgA normally rise in starving rods engaged in mound building (3), but the addition of sufficient nutrients triggers rapid proteolysis of MrpC and prevents the normal increase in transcripts for proteins involved in sporulation (38, 39). Combinatorial control of gene expression by MrpC and FruA appears to ensure that only starving rods (in which MrpC and FruA are synthesized and stably accumulate) aligned in mounds (which enhances short-range C-signaling between rods, presumably activating FruA within them) commit to spore formation (7, 19, 31, 38, 46).

Among the genes combinatorially controlled by MrpC and FruA, early studies showed that mutations in the *dev* operon can cause severe defects in sporulation (47–50). The *dev* operon includes eight genes comprising a CRISPR-Cas system that has been proposed to protect developing *M. xanthus* against bacteriophage lysogenization (51). The first gene in the *dev* operon, *devI*, codes for a 40-residue protein that strongly inhibits sporulation when overproduced (52), delays sporulation by about 6 h when produced at the normal level (29, 53), and exerts weak positive autoregulation on *dev* transcript accumulation (29). In contrast, in-frame deletions in three genes of the *dev* operon (*devTRS*) increase accumulation of the *dev* transcript 10-fold during development (29, 52, 53), indicating that DevTRS proteins exert strong negative autoregulation (Fig. 1). The negative autoregulation by DevTRS proteins is crucial in order to prevent DevI overproduction from inhibiting sporulation (52, 53) (Fig. 1). The DevTRS proteins are similar to Cas8a1, Cas7, and Cas5, respectively, of the <u>C</u>RISPR-<u>as</u>sociated <u>c</u>omplex for <u>a</u>ntiviral <u>de</u>fense (Cascade) (51–54).

The products of several other operons act late during the developmental process, impacting the metabolism of nascent spores (*fadIJ*) and the biogenesis of their coat polysaccharide (*exoA-I*, *exoL-P*, and *nfsA-H*), but the regulation of these operons is not fully understood (Fig. 1). The putative *fadIJ* operon is normally induced twofold in sporulating cells, but not in *csgA*, *fruA*, or *mrpC* mutants (55). Recently published data support that *fadIJ* are co-transcribed as an operon FadIJ are enzymes in a fatty acid β-oxidation pathway that can impact spore structure and resistance properties (55). The *nfsA-H* operon is up-regulated in spores formed during the starvation-induced developmental process and in spores induced artificially by glycerol addition The starvation-induced up-regulation fails in a *csgA* mutant, is diminished in a *devRS* mutant, and, unusually, is greater in a *fruA* mutant, indicative of negative regulation by FruA (57). The NfsA-H proteins form a complex involved in biogenesis of the coat polysaccharide to generate compact and rigid stress-resistant spores (58–60). The *exoA-I* operon (58) is normally induced late in development, but not in *csgA* or *devRS* mutants (61). Transcription depends on binding of FruA (62) and Nla6 (63) to the *exoA-I* promoter region. Interestingly, Nla6 is part of a cascade of enhancer-binding proteins (EBPs) that responds to starvation early in development (32) and leads to production of MrpC and FruA (7) (Fig. 1). The Nla6 DNA-binding domain (DBD) also binds to the promoter region of the predicted *MXAN_3259*–*MXAN_3263* operon and appears to directly regulate transcription (63). Recently published data support that the five genes are co-transcribed as an operon (56), which was renamed the *exoL-P* operon based on functional studies (64). The ExoA-I and ExoL-P proteins are involved in the synthesis, export, and modification of the spore coat polysaccharide (58, 59, 64).

Here, we report systematic investigation of the late-acting *fadIJ*, *nfsA-H*, *exoA-I*, and *exoL-P* operons with respect to the phenotypes of mutants and the regulation of gene expression. We found that the *nfs* and *exo* mutants exhibit defects in sporulation earlier than established previously, whereas a *fad* mutant initially made more sonication-resistant spores than a “wild-type” (WT) strain, but later made a similar number of mature spores. Cooperative binding of MrpC and C-signal-activated FruA appears to explain up-regulation of *fadIJ* late in development, but not the regulation of *nfsA-H*, *exoA-I*, and *exoL-P.* FruA negatively regulates all four operons early in development and MrpC affects each operon differently. Nla6 up-regulates all four operons and we discovered that its DBD binds cooperatively with FruA to the *nfsA-H*, *exoL-P*, and *exoA-I* promoter regions, and also binds cooperatively with MrpC to the *nfsA-H* promoter region. We also found that all four operons are subject to down-regulation by a CRISPR-Cas system, especially when the DevI component is overexpressed, as in a *devS* mutant. We conclude that three transcription factors and a CRISPR-Cas component regulate the late-acting operons and we propose that complex, differential regulation of genes involved in spore metabolism and coat biogenesis produces spores suited to withstand environmental insults.

## Results

### *nfs* and *exo* mutants exhibit defects early in the sporulation process

In previous work, we established methods to systematically analyze *M. xanthus* development under submerged culture conditions (29). Our method of submerged culture involves growing cells in nutrient medium followed by sedimentation, resuspension in starvation buffer, and incubation in a plastic container. Cells attach to the container bottom and develop under a layer of buffer.

To determine the effects of mutations in the late-acting operons under our conditions of submerged culture, we compared the development of mutants with that of WT laboratory strain DK1622 (65) from which the mutant strains were derived. Specifically, we examined the effects of an insertion in *fadI* (55), a deletion of the entire *nfsA-H* operon (57), an insertion in *exoC* (58, 61), and an insertion in *exoL* (63, 64). As expected (29), the WT strain formed mounds by 18 h poststarvation (PS) and the mounds began to darken by 30 h (Fig. 2). Darkening typically correlates with spore formation. The *fadI* mutant was indistinguishable from the WT strain. The *nfsA-H* mutant was delayed by about 3 h in mound formation and appeared to be delayed and reduced in mound darkening. The *exoC* and *exoL* mutants formed normal-looking mounds by 18 h, but subsequent mound darkening was reduced compared to the WT strain.

**Figure 2.**
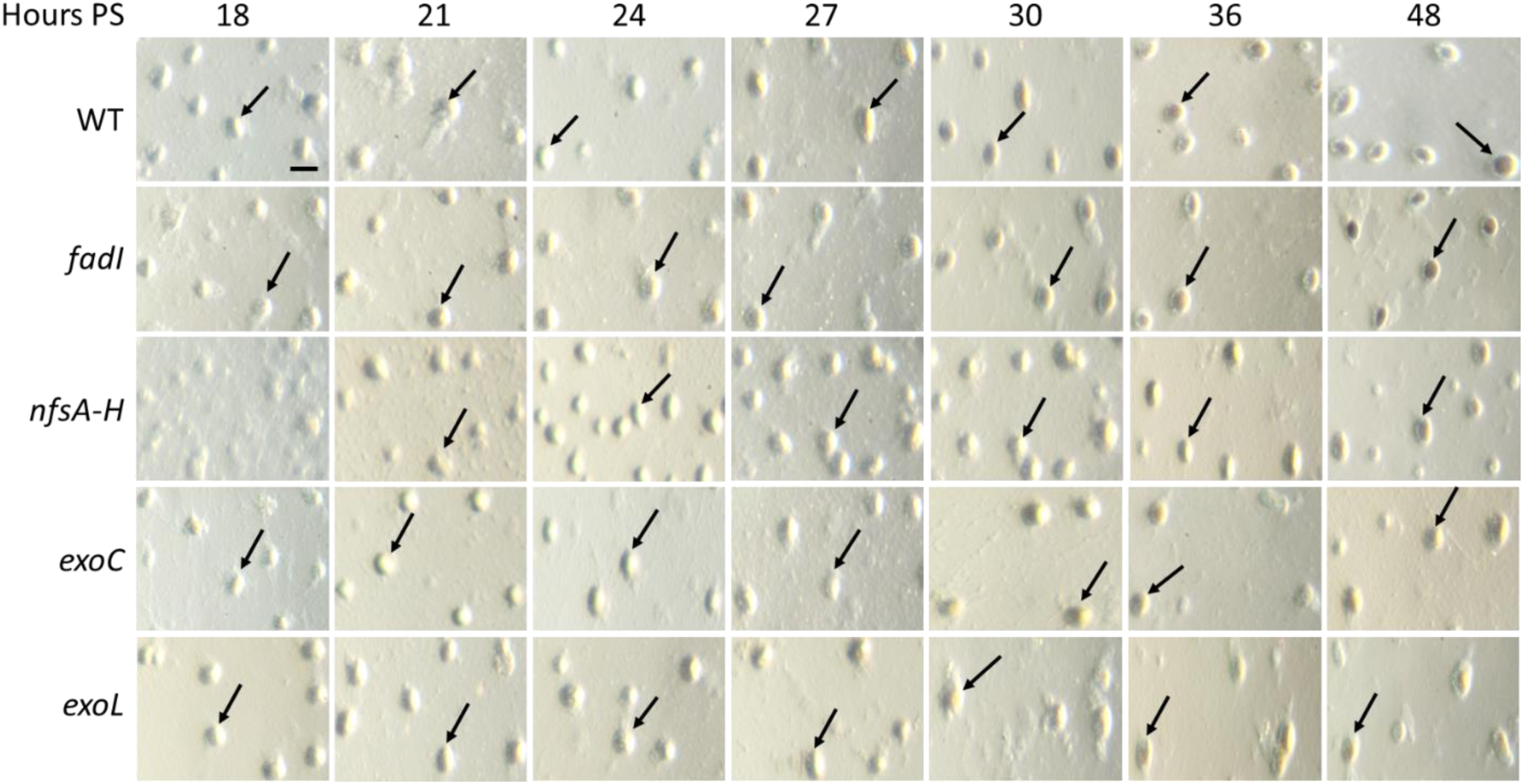
Development of *M. xanthus* strains. The WT strain and its indicated mutant derivatives were subjected to starvation under submerged culture condition and microscopic images were obtained at the indicated times PS. The WT strain and all the mutants except *nfsA-H* formed compact mounds by 18 h (an arrow points to one in each panel). The *nfsA-H* mutant formed compact mounds by 21 h. Mounds of the WT strain began to darken by 30 h. Mounds of the *nfsA-H*, *exoC*, and *exoL* mutants darkened by 30 h but remained slightly less dark than mounds of the WT strain and the *fadI* mutant by 48 h. Bar, 100 μm. Similar results were observed in at least three biological replicates.

To quantify changes at the cellular level, we harvested samples from submerged culture and either treated the samples with glutaraldehyde to fix cells or left the samples untreated (29). Using the untreated samples, we quantified “sonication-resistant spores” microscopically and “mature spores” that were heat- and sonication-resistant and capable of germination and colony formation on nutrient agar. We used the fixed samples to quantify “sonication-sensitive cells” microscopically (i.e., the total number of cells observed in the fixed sample minus the number of sonication-resistant spores observed in the corresponding untreated sample). The vast majority of sonication-sensitive cells were rod-shaped, but we also observed a small percentage of “transitioning cells” (TCs) intermediate in morphology between rods and spores (46) in some of the fixed samples.

As expected (29), we first observed sonication-resistant spores at 27 h PS in samples of the WT strain, and the spores rose to ∼2% at 48 h (Fig. 3). The *fadI* mutant formed more spores than the WT strain. Consistent with the observed mound darkening defects (Fig. 2), the *nfsA-H* mutant formed less spores than the WT strain, and the *exoC* and *exoL* mutants did not form sonication-resistant spores (at a detection limit of 0.05%) (Fig. 3 and Table S1). Neither did the *exoC* and *exoL* mutants form mature spores at 72 h (at a detection limit of 0.00001%), while the *nfsA-H* mutant made ∼15% as many mature spores as the WT strain, and the *fadI* mutant made a similar number of mature spores as the WT strain (Table S1).

**Figure 3.**
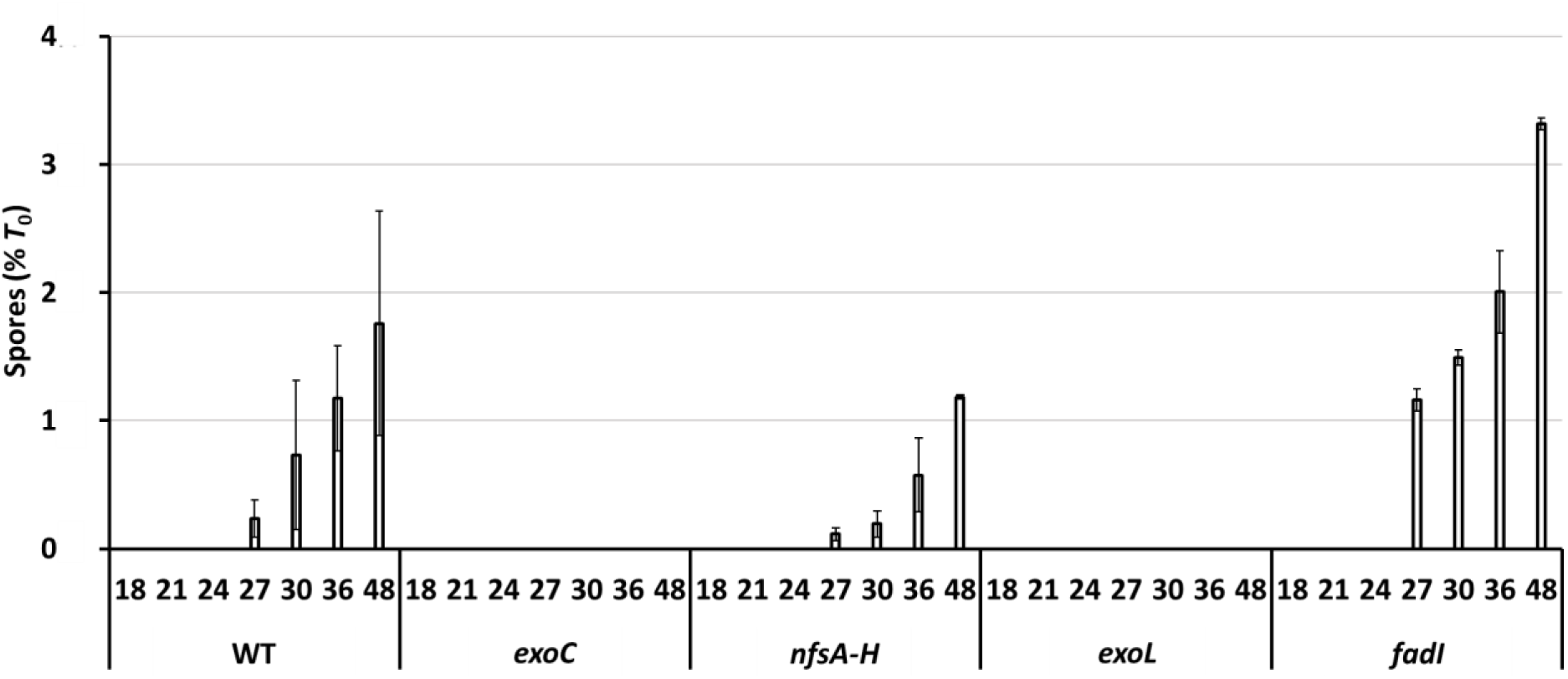
Sonication-resistant spores during *M. xanthus* development. The WT strain and its indicated mutant derivatives were subjected to starvation under submerged culture conditions. Samples were collected at the indicated times PS for quantification of sonication-resistant spores. Values are expressed as a percentage of the number of rod-shaped cells present at the time starvation initiated development (*T*_0_) (Table S1). Bars show the average of three biological replicates and error bars indicate one standard deviation.

The WT strain exhibited a decline of sonication-sensitive cells during development similar to that reported previously (29), with only ∼30% of the cells remaining at 18 h PS and only ∼6% remaining at 48 h (Fig. S1A), presumably reflecting lysis of the majority of cells during submerged culture development (3). The mutants showed similar decreases in cell number as the WT strain (Fig. S1A). As noted above, we observed a small percentage of TCs in some of the fixed samples. For the WT strain, ∼3% of the cells were TCs at 24 h (Fig. S1B), when ∼20% of the cells remained (Fig. S1A). Thus, TCs comprised 15% of the total population at 24 h, before sonication-resistant spores were observed (Fig. 3), in agreement with *in situ* observations using confocal microscopy (46). The *fadI* mutant showed about half as many TCs as the WT strain at most time points (Fig. S1B), perhaps due to the *fadI* mutant forming more sonication-resistant spores (Fig. 3). The *exo* and *nfs* mutants exhibited less TCs at 24 h (∼0.4-0.7%), but in each case the percentage rose to at least 1% later (Fig. S1B), suggesting that these mutants begin to change shape, but are impaired early in the process of making spores.

We conclude that the *exoC* and *exoL* mutants showed stronger defects in sporulation than the *nfsA-H* mutant, whereas the *fadI* mutant initially made more sonication-resistant spores than the WT strain, but later made a similar number of mature spores. All four mutants exhibited less TCs than the WT strain early in the sporulation process.

### Transcript levels from the late-acting operons are very low in the absence of C-signaling

To determine the effects of loss of C-signaling on expression of the late-acting operons, we used methods established previously (29) to measure transcript levels in the WT strain and a *csgA* insertion mutant at 18-30 h PS, the period leading up to and including the time that many cells commit to spore formation and sonication-resistant spores begin to be observed (38) (Fig. 3). As expected for the WT strain (39), transcript levels rose during development (Fig. 4). As noted previously (53), the *exoA* transcript level varied greatly between biological replicates (Fig. 4A), which we do not understand. The *exoL* transcript level likewise varied greatly (Fig. 4B). The *nfsA* (Fig. 4C) and *fadI* (Fig. 4D) transcript levels varied less and on average increased only ∼fourfold, much less than the *exoA* (∼24-fold) and *exoL* (∼70-fold) transcript levels.

**Figure 4.**
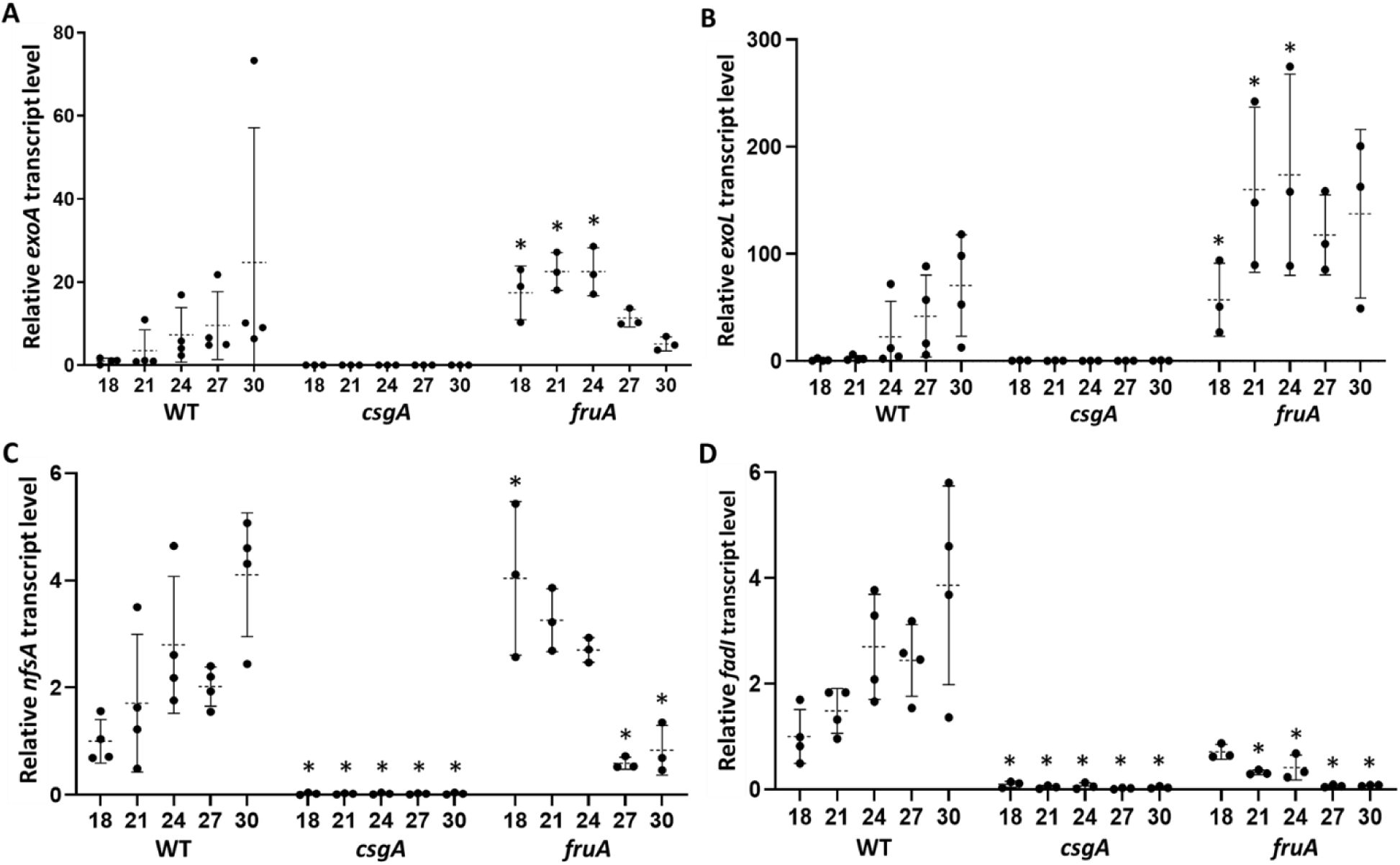
Transcript levels in the WT strain and the *csgA* and *fruA* mutants late in *M. xanthus* development. The WT strain and its mutant derivatives were subjected to starvation under submerged culture conditions. Samples were collected at the indicated times PS for measurement of the *exoA* (A), *exoL* (B), *nfsA* (C), and *fadI* (D) transcript levels by RT-qPCR. Graphs show the data points and the average of at least three biological replicates, relative to the WT strain at 18 h PS, and error bars indicate one standard deviation. Asterisks indicate a difference (*p* < 0.05 in Student’s two-tailed *t*-tests) from the WT strain at the corresponding time PS.

In the *csgA* mutant, the transcript levels did not increase during development (Fig. 4, asterisks indicate *p* < 0.05 in Student’s two-tailed *t*-tests comparing the mutant to the WT strain at each time point). Asterisks are absent above the *exoA* and *exoL* transcript levels of the *csgA* mutant because the statistical test yielded *p* > 0.05 due to the large variation between biological replicates of the WT strain. Nevertheless, the *exoA* and *exoL* transcript levels of the *csgA* mutant were low for all replicates at each time point. The low transcript levels could be due to decreased synthesis and/or increased degradation in the *csgA* mutant compared with the WT strain. To measure the transcript degradation rates, we added rifampicin to inhibit transcription at 30 h PS and determined the transcript levels at intervals thereafter. The transcript degradation rates did not differ significantly between the *csgA* mutant and the WT strain (Fig. 5), suggesting that decreased synthesis primarily accounts for the low transcript levels in the absence of C-signaling. In agreement, reporter activity from fusions to *exoC* (61), *nfsA* (57), and *fadI* (55) was very low in *csgA* mutants relative to WT strains during development.

**Figure 5.**
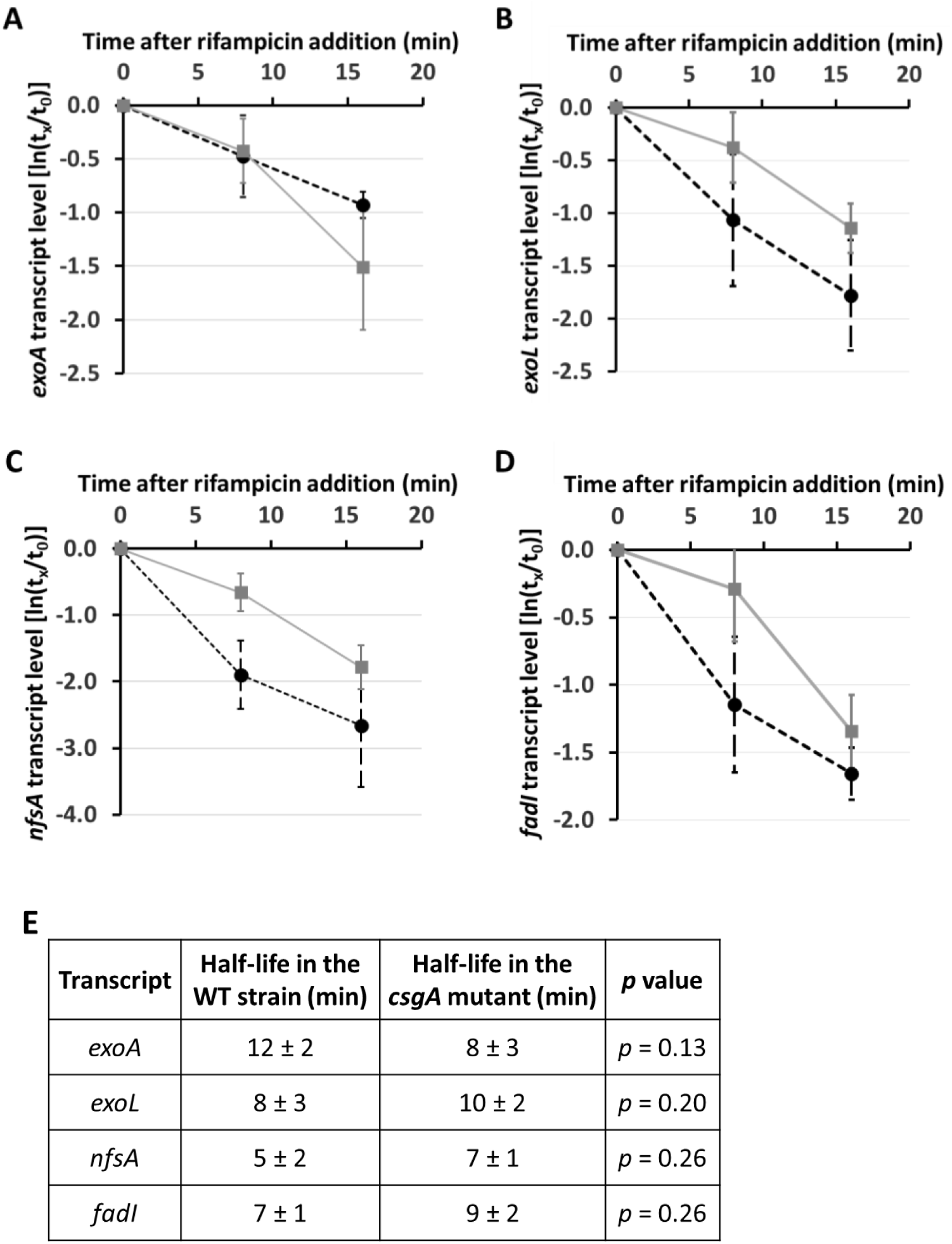
Transcript stability in the WT strain and the *csgA* mutant late in *M. xanthus* development. The WT strain and the *csgA* mutant were subjected to starvation under submerged culture conditions for 30 h. The overlay was replaced with fresh starvation buffer containing rifampicin (50 μg/mL) and samples were collected immediately (*t*_0_) and at the indicated times (*t*_x_) for measurement of the *exoA* (A), *exoL* (B), *nfsA* (C), and *fadI* (D) transcript levels by RT-qPCR. Transcript levels at *t*_x_ were normalized to that at *t*_0_ for each of three biological replicates and used to determine the transcript half-life for each replicate. The graph shows the average ln(*t*_x_/*t*_0_) and one standard deviation for the three biological replicates of the WT strain (black dashed line) and the *csgA* mutant (gray solid line). The average half-life and one standard deviation, as well as the *p* value from a Student’s two-tailed *t*-test, are reported in (E).

Considerable evidence supports a model in which C-signaling activates FruA posttranslationally in order to increase transcription of genes during *M. xanthus* development (28, 29). We showed previously that the FruA level is about twofold lower in the *csgA* mutant than in the WT strain at 18-30 h PS (29). Boosting the FruA level in the *csgA* mutant to the WT level using a vanillate-inducible promoter (P*_van_*) fused to *fruA* did not increase the transcript levels of five genes or operons (*dev*, *fmgA*, *fmgBC*, *fmgD*, and *fmgE*) known to be under combinatorial control of FruA and MrpC. Neither did boosting the level of FruA D59E (with a phosphomimetic substitution in its receiver domain) using a P*_van_-fruA D59E* fusion.

We tested whether boosting the level of FruA or its D59E variant in the *csgA* mutant increases the transcript levels of the late-acting operons. The transcript levels remained low in all cases (Fig. S2), suggesting that neither a low level of FruA nor a lack of D59 phosphorylation causes the transcript levels to remain low in the *csgA* mutant. Rather, these results are consistent with evidence that C-signaling regulates FruA activity by a mechanism other than phosphorylation of its receiver domain (29, 31).

### Transcript levels from the late-acting operons are unexpectedly high in a *fruA* mutant

If C-signaling activates FruA and the activated FruA increases transcription of the late-acting operons, then transcript levels are expected to be low in a *fruA* insertion mutant, as observed for the *dev* operon (29). Instead, the *exoA* and *exoL* transcript levels were greater in the *fruA* mutant than in the WT strain at 18 to 24 h PS (Fig. 4A and 4B), and the *nfsA* transcript level in the *fruA* mutant exceeded that in the WT strain at 18 h (Fig. 4C), suggesting that FruA is a negative regulator of some late-acting operons early in development. In contrast, the *fadI* transcript level in the *fruA* mutant was similar to that in the WT strain at 18 h and less than that in the WT strain at later times (Fig. 4D), consistent with a model in which C-signaling activates FruA and the activated FruA increases *fadI* transcription. Activated FruA may also increase *nfsA* transcription at 27 and 30 h, since the *nfsA* transcript level in the *fruA* mutant was less than that in the WT strain at those times (Fig. 4C).

Interestingly, the average transcript levels from the late-acting operons were greater in the *fruA* mutant than in the *csgA* mutant, except for the very low *fadI* transcript level in both mutants at 27 and 30 h PS (Fig. 4). As noted above, FruA is present, but its level is about twofold lower in the *csgA* mutant than in the WT strain at 18-30 h (29). In the *csgA* mutant, FruA is expected to remain unactivated due to the absence of C-signaling (28, 29). Therefore, it is possible that unactivated FruA negatively regulates the late-acting operons in the *csgA* mutant. Negative regulation by unactivated FruA may also account for the lower levels of *exoA*, *exoL*, and *nfsA* transcripts we observed early in development of the WT strain as compared with the *fruA* mutant (Fig. 4) (see Discussion).

### The exoA, exoL, and nfsA transcript levels of mrpC and fruA mutants differ

MrpC appears to directly stimulate transcription from the *fruA* promoter (33). In agreement, FruA was not detected in an *mrpC* in-frame deletion mutant at 18-30 h PS (29). Hence, the *mrpC* mutant lacks both MrpC and a detectable level of FruA. To compare the *mrpC* mutant with the *fruA* mutant (lacking FruA but with MrpC present at its normal level) (29), we measured the transcript levels from the late-acting operons in the mutants and the WT strain in parallel. Strikingly, the *exoA*, *exoL*, and *nfsA* transcript levels in the *mrpC* and *fruA* mutants differed, and in each case the pattern of effects was unique. The average *exoA* transcript level was greater in the *mrpC* mutant than in the *fruA* mutant or in the WT strain at all times (Fig. 6A, asterisks above brackets indicate *p* < 0.05 in Student’s two-tailed *t*-tests comparing the mutants at the same time point and asterisks above error bars compare a mutant to the WT strain at the same time point). The comparison of mutants suggests that MrpC negatively regulates the *exoA* transcript level independently of FruA, since the loss of both transcription factors in the *mrpC* mutant increased the *exoA* transcript level more than loss of only FruA in the *fruA* mutant. The *exoL* transcript level in the *mrpC* mutant exceeded that in the WT strain at 18 and 24 h, and on average exceeded that in the *fruA* mutant at 18 h, but on average was less than in the *fruA* mutant at 24 h, and at 30 h the difference met a test of statistical significance (Fig. 6B). In this case, comparison of the mutants suggests that, independently of FruA, MrpC negatively regulates the *exoL* transcript level early in development and positively regulates it later. The average *nfsA* transcript level was lower in the *mrpC* mutant than in the *fruA* mutant or in the WT strain at all times (Fig. 6C), suggesting that MrpC positively regulates the *nfsA* transcript level independently of FruA (i.e., opposite the regulation of *exoA*). The *fadI* transcript level was lower in both mutants than in the WT strain at all times (Fig. 6D), consistent with positive combinatorial control by MrpC and FruA, as observed for *dev* (42), *fmg* (31, 43–45), and likely many other developmentally-regulated genes or operons (41).

**Figure 6.**
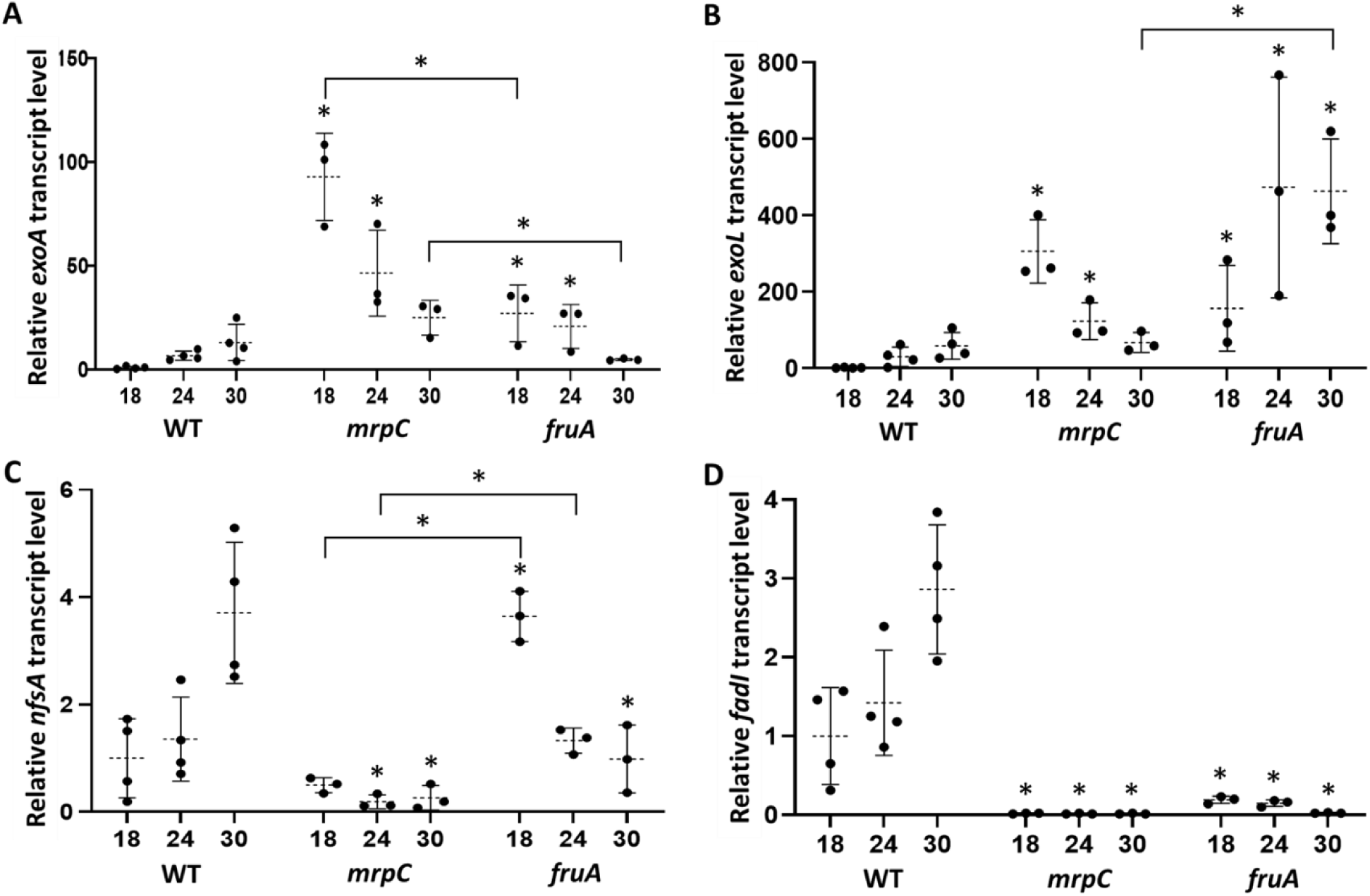
Transcript levels in the WT strain and the *mrpC* and *fruA* mutants late in *M. xanthus* development. The WT strain and its mutant derivatives were subjected to starvation under submerged culture conditions. Samples were collected at the indicated times PS for measurement of *exoA* (A), *exoL* (B), *nfsA* (C), and *fadI* (D) transcript levels by RT-qPCR. Graphs show the data points and the average of at least three biological replicates, relative to the WT strain at 18 h PS, and error bars indicate one standard deviation. Asterisks above error bars indicate a difference (*p* < 0.05 in Student’s two-tailed *t*-tests) from the WT strain at the corresponding time PS and asterisks above brackets indicate a difference between the mutants.

To determine whether the lack of both MrpC and a detectable level of FruA in the *mrpC* mutant affects the transcript degradation rates, we added rifampicin to inhibit transcription at 18 h PS and determined the transcript levels at intervals thereafter. We chose 18 h for this analysis since the *exoA* (Fig. 6A) and *exoL* (Fig. 6B) transcript levels were elevated in the *mrpC* mutant relative to the WT strain at that time. The transcript degradation rates did not differ significantly between the *mrpC* mutant and the WT strain (Fig. S3), suggesting that increased synthesis (rather than decreased degradation) primarily accounts for the elevated *exoA* and *exoL* transcript levels in the *mrpC* mutant.

### FruA can positively regulate *exoA*, *exoL*, and *nfsA* in the absence of MrpC

To examine the effects of FruA in the absence of MrpC, we used the P*_van_-fruA* fusion mentioned previously (29) (Fig. S2) to produce FruA in the *mrpC* mutant. The inducer (vanillate) was added during growth and at 0 h PS. At 6 h, the FruA level in the *mrpC* P*_van_-fruA* mutant was about threefold greater than in the WT strain, but their FruA level was similar at 12 and 18 h (Fig. S4A). Interestingly, the *mrpC* P*_van_-fruA* mutant formed immature mounds by 12 h (Fig. S5), although subsequent mound darkening was reduced compared to the WT strain (Fig. S6) and spores were not detected (Table S1). Nevertheless, mound formation suggests that C-signaling is activating FruA to some extent, since a complete block, as in *mrpC*, *fruA*, and *csgA* mutants, results in no mound formation (29) (Fig. S5).

To determine the effects of FruA on regulation of the late-acting operons in the absence of MrpC, we measured the transcript levels in the *mrpC* P*_van_-fruA* mutant in parallel with those in the *mrpC* and *fruA* mutants and the WT strain at 6-18 h PS. We chose to collect samples during that period of development since FruA was present (Fig. S4A) and appeared to be active (Fig. S5) in the *mrpC* P*_van_-fruA* mutant and the WT strain, as described above. The *exoA* (Fig. 7A) and *exoL* (Fig. 7B) transcript levels in the *mrpC* P*_van_-fruA* mutant exceeded those in the WT strain at all times, and on average exceeded those in the *mrpC* and *fruA* mutants, except the *exoL* transcript level in the *mrpC* mutant was similar to that in the *mrpC* P*_van_-fruA* mutant at 18 h.

**Figure 7.**
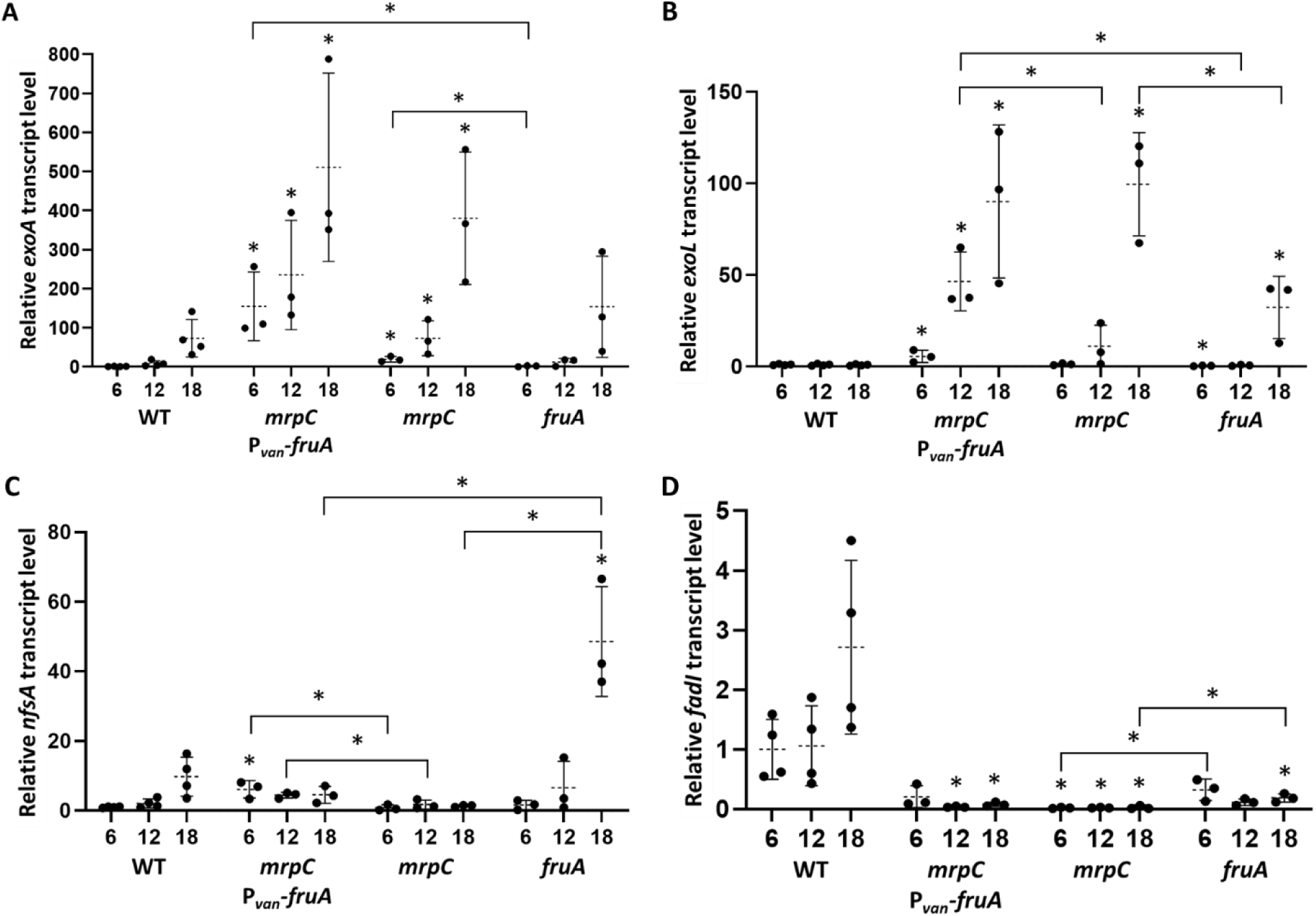
Transcript levels in the WT strain, the *mrpC* mutant with inducible *fruA*, and the *mrpC* and *fruA* mutants early in *M. xanthus* development. The WT strain and its mutant derivatives were subjected to starvation under submerged culture conditions. Samples were collected at the indicated times PS for measurement of *exoA* (A), *exoL* (B), *nfsA* (C), and *fadI* (D) transcript levels by RT-qPCR. Expression of *fruA* fused to P*_van_* was induced with vanillate (0.5 mM) during growth and at 0 h PS. Graphs show the data points and the average of at least three biological replicates, relative to the WT strain at 6 h PS, and error bars indicate one standard deviation. Asterisks above error bars indicate a difference (*p* < 0.05 in Student’s two-tailed *t*-tests) from the WT strain at the corresponding time PS and asterisks above brackets indicate a difference between the mutants.

We conclude that FruA can positively regulate the *exoA* and *exoL* transcript levels in the absence of MrpC. This regulation may involve C-signal-activated FruA since the *mrpC* P*_van_-fruA* mutant formed immature mounds (Fig. S5 and S6), whereas *mrpC*, *fruA*, and *csgA* mutants fail to form mounds (29) (Fig. S5). The *nfsA* transcript level in the *mrpC* P*_van_-fruA* strain exceeded that in the WT strain at 6 h and that in the *mrpC* mutant at 6 and 12 h (Fig. 7C), also indicative of positive regulation by FruA (perhaps activated FruA) in the absence of MrpC. On the other hand, the *fadI* transcript level in the *mrpC* P*_van_-fruA* strain was less than that in the WT strain (Fig. 7D), as observed for the *mrpC*, *fruA*, and *csgA* mutants (Fig. 4D, 6D, and 7D), supporting that positive regulation requires both MrpC and activated FruA.

### Nla6 positively regulates the transcript levels of the late-acting operons

The Nla6 transcription factor appears to be a direct regulator of the *exoA-I* and *exoL-P* operons since a protein with the *E. coli* maltose-binding protein (MBP) fused to the Nla6 DBD (MBP-Nla6 DBD) bound to promoter region DNA fragments (63). A comparison of the *exoA* and *exoL* transcript levels in an *nla6* insertion mutant and a WT strain suggested that Nla6 positively regulates both during the first 8 h PS, and negatively regulates both at 24 h (63). To facilitate planned strain construction, we constructed a new *nla6* insertion mutant and compared it with the one characterized previously (63, 66). The new mutant is tetracycline-resistant (Tc^r^) and the one described previously is kanamycin-resistant (Km^r^). Both mutants formed immature mounds by 12 h, but failed to progress to more mature mounds with distinct, round edges by 18 h (Fig. S5). Later during development, the Km^r^ *nla6* mutant mounds matured somewhat at 24-30 h, but failed to darken by 36-48 h (Fig. S6). The Tc^r^ *nla6* mutant mounds did not mature until 36 h and also failed to darken by 48 h. Spores were not detected for either mutant (Table S1).

The two *nla6* mutants were indistinguishable in terms of the molecular markers we tested. Since our results showed that FruA and MrpC impact transcript levels of the late-acting operons at 18 h PS (Fig. 4 and 6) and in some cases earlier during development (Fig. 7), we measured the levels of FruA, MrpC, and transcripts from the late-acting operons in the *nla6* mutants and the WT strain at 6-18 h. The FruA and MrpC protein levels in the *nla6* mutants were similar to those in the WT strain at 6-18 h (Fig. S4). The *exoA*, *nfsA*, and *fadI* transcript levels in the WT strain increased at 18 h, but those in the *nla6* mutants failed to increase (Fig. 8), suggesting that Nla6 positively regulates those transcript levels. The *exoL* transcript level in the WT strain did not increase at 18 h, but on average that in the *nla6* mutants was less (Fig. 8B), suggesting that Nla6 also positively regulates the *exoL* transcript level. Because the *exoA*, *exoL*, and *nfsA* transcript levels of the *mrpC* and/or *fruA* mutants exceeded those of the WT strain early in development (Fig. 4, 6, and 7), we tried to construct *mrpC nla6* and *fruA nla6* double mutants, but our efforts were unsuccessful, so we were unable to determine whether positive regulation by Nla6 could account for the elevated transcript levels in the *mrpC* and *fruA* mutants relative to the WT strain.

**Figure 8.**
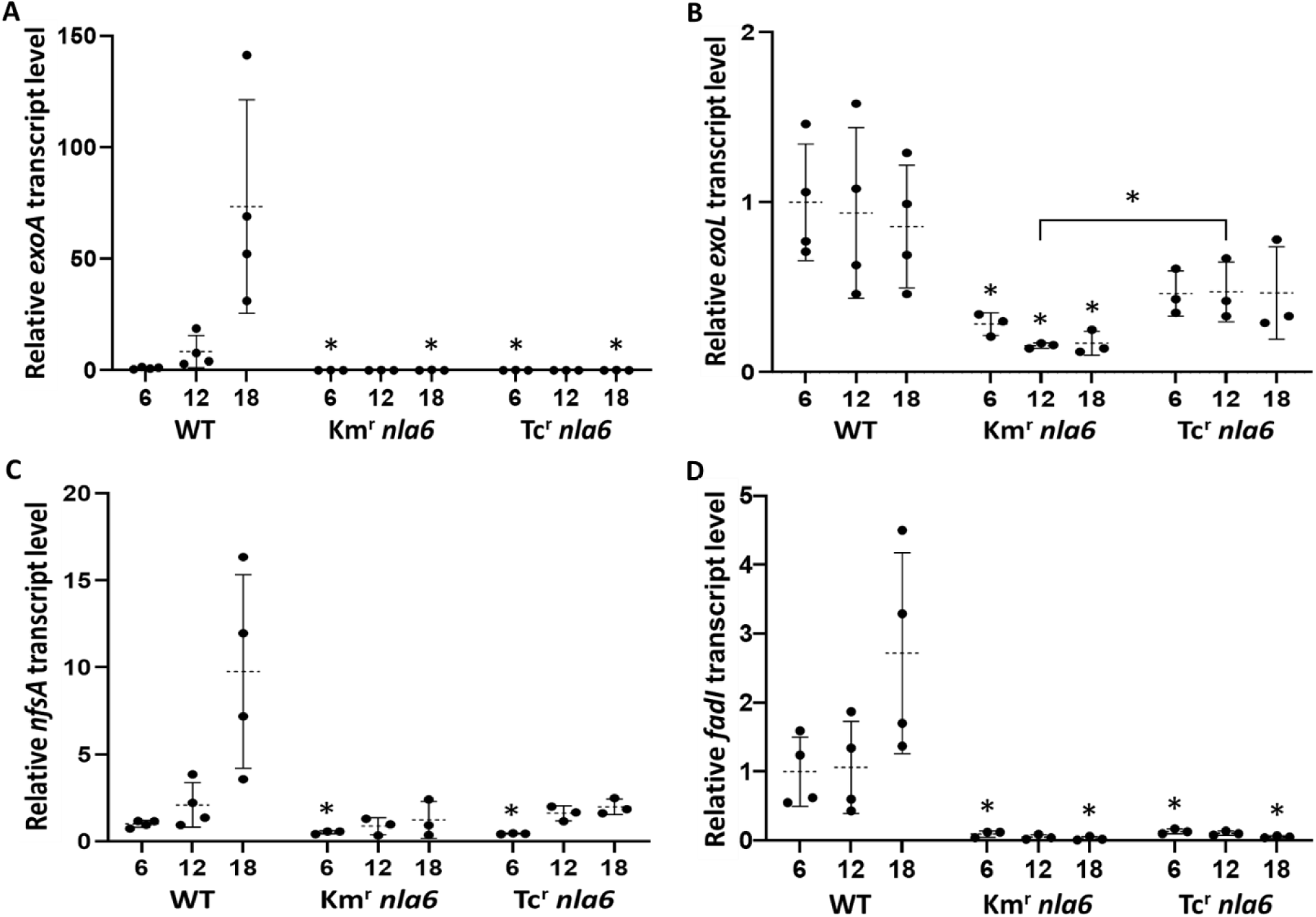
Transcript levels in *nla6* mutants early in *M. xanthus* development. The WT strain and its mutant derivatives were subjected to starvation under submerged culture conditions. Samples were collected at the indicated times PS for measurement of *exoA* (A), *exoL* (B), *nfsA* (C), and *fadI* (D) transcript levels by RT-qPCR. Graphs show the data points and the average of at least three biological replicates, relative to the WT strain at 6 h PS, and error bars indicate one standard deviation. Asterisks indicate a difference (*p* < 0.05 in Student’s two-tailed *t*-tests) from the WT strain at the corresponding time PS and the asterisk above the bracket indicates a difference between the mutants.

### Binding of MBP-Nla6 DBD, MrpC, and FruA differs at the promoter region of each of the late-acting operons

As a possible indication of direct regulation by Nla6, MrpC, and FruA, we tested for binding to a DNA fragment upstream of the predicted start codon of the first gene in each of the late-acting operons. Figure S7 shows features of each promoter region fragment based on previous work. FruA and MrpC were shown to bind cooperatively to the *dev* promoter region fragment (42) (Fig. S7A), which was used here as a control. The FruA DBD was shown to bind to at least two sites in the *exoA-I* promoter region fragment (62) and MBP-Nla6 DBD was also shown to bind, presumably *via* interaction with DNA sequences similar to a half-site consensus sequence (63) (Fig. S7B). MBP-Nla6 DBD was also shown to bind to the *exoL-P* promoter region fragment (63) and three matches to the half-site consensus sequence were identified (Fig. S7C). ChIP-seq analysis suggested that MrpC binds to a site upstream of *nfsA-H* at 18 h PS (41) (Fig. S7D). Here, we used electrophoretic mobility shift assays (EMSAs) to test for binding of purified MBP-Nla6 DBD, MrpC, and FruA separately, and in combination because MrpC and FruA were shown previously to bind cooperatively to the promoter regions of *dev* (42), *fmg* (31, 43–45), and other developmentally-regulated genes or operons (41).

As expected (42), MrpC and FruA bound individually to the *dev* promoter region fragment (Fig. 9A, lanes 4, 5, 7, 8), and the combination of MrpC and FruA exhibited greater than additive binding (lanes 11, 14), indicative of cooperative binding. MrpC and FruA DBD also appeared to bind cooperatively (Fig. S8A, lane 4), indicating the N-terminal domain of FruA was not required. Binding of MBP-Nla6 DBD was not detected at 4 μM (Fig. 9A, lane 2), consistent with previous reports (32, 63), but binding was detected at 16 μM and increased at greater concentrations (Fig. S8A, lanes 8-10).

**Figure 9.**
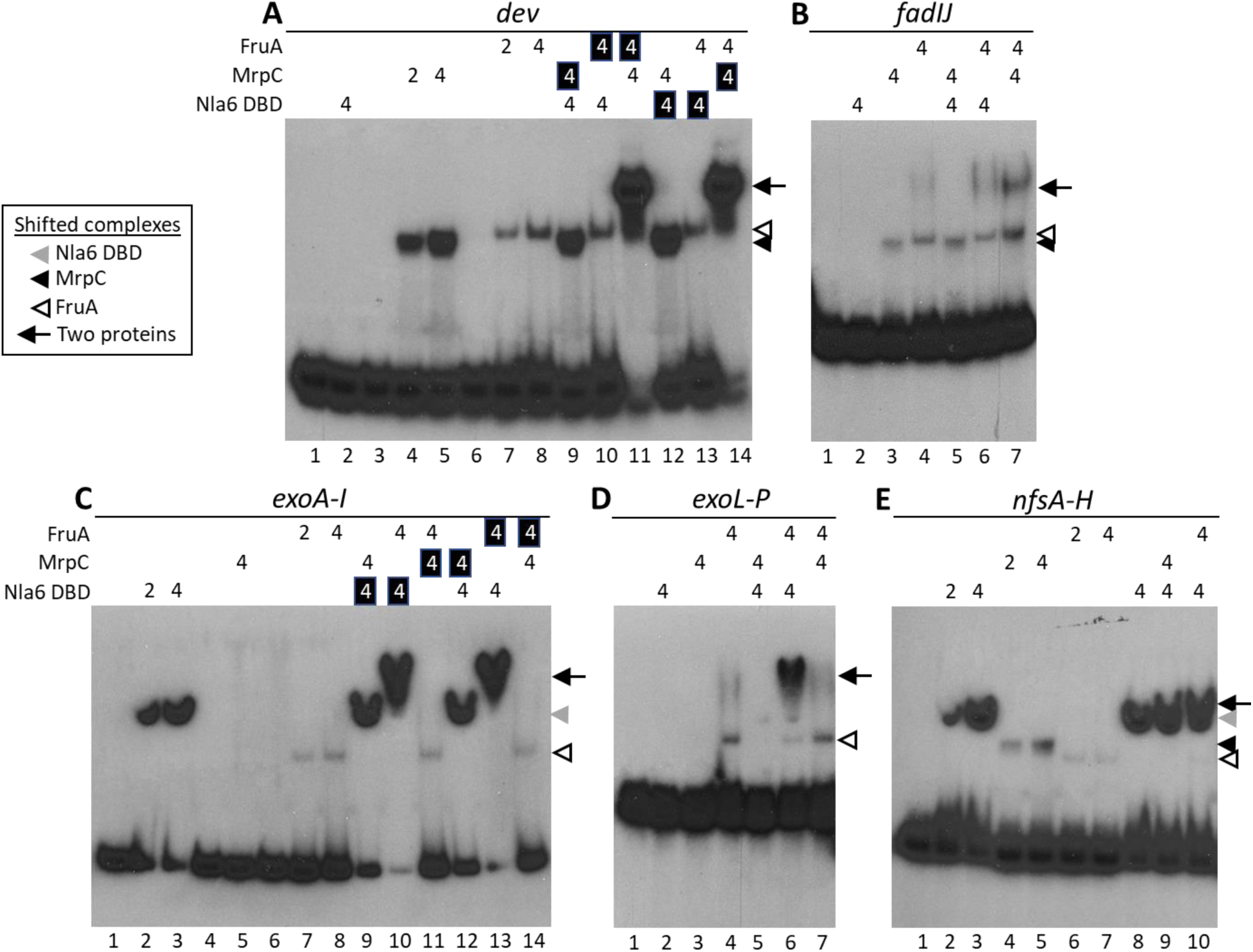
Binding of the Nla6 DBD, MrpC, and FruA to the promoter regions of the late-acting operons. EMSAs were performed with ^32^P-labeled DNA fragments (2 nM) of the *dev* (−114 to - 19) (A), *fadIJ* (−214 to −5) (B), *exoA-I* (−171 to −1) (C), *exoL-P* (−217 to +56) (D), and *nfsA-H* (−166 to +35) (E) promoter regions, and MBP-Nla6 DBD, His_6_-MrpC, and FruA-His_6_ at the μM concentrations indicated by numbers above each autoradiographic image. A white number on a black background indicates a protein that was added to the DNA-binding reaction 10 min later than the other protein. The positions of migration of shifted complexes are indicated along the side of each image by arrowheads for binding of individual proteins and by arrows for binding of two proteins (see boxed key). Lanes are numbered below images.

The *fadIJ* promoter region fragment yielded results that were in some ways similar to those for the *dev* fragment. MrpC and FruA bound individually to the *fadIJ* fragment (Fig. 9B, lanes 3, 4), and the combination of MrpC and FruA appeared to bind cooperatively (lane 7), but with much less cooperative binding than to the *dev* fragment. The order of MrpC and FruA addition did not affect cooperative binding to either fragment (Fig. 9A, lanes 11, 14 and Fig. S8B, lanes 5, 6). However, MrpC and FruA DBD did not bind cooperatively to the *fadIJ* fragment (Fig. S8B, lane 10), indicating a requirement for the FruA N-terminal domain, in contrast to the *dev* fragment. On the other hand, for both fragments, MBP-Nla6 DBD binding was not detected at 4 μM and at that concentration did not affect binding of MrpC or FruA (Fig. 9A, lanes 2, 9, 10, 12, 13 and Fig. 9B, lanes 2, 5, 6). Also like the *dev* fragment, MBP-Nla6 DBD binding to the *fadIJ* fragment was detected at 16 μM and increased at higher concentrations (Fig. S8B, lanes 14-16). The apparent cooperative binding of MrpC and FruA to the *fadIJ* fragment, together with our measurements of the *fadI* transcript level, support a similar mechanism of developmental regulation of the *fadIJ* operon as proposed for the *dev* operon (29, 42), the *fmg* genes (31, 43–45), and likely many other genes (41), in which C-signaling activates FruA to bind DNA cooperatively with MrpC and increase transcription.

MBP-Nla6 DBD at 2 μM bound to the *exoA-I* promoter region fragment (Fig. 9C, lane 2), as expected (63). FruA also bound (Fig. 9C, lanes 7, 8), as expected based on FruA DBD binding observed previously (62). Strikingly, MBP-Nla6 DBD and FruA appeared to bind cooperatively and the order of addition made no difference (Fig. 9C, lanes 10, 13 and Fig. S8C, lanes 4-6). FruA DBD exhibited very little cooperative binding with MBP-Nla6 DBD (Fig. S8C, lane 10), suggesting the N-terminal domain of FruA plays an important role. Although MrpC appeared to negatively regulate the *exoA* transcript level independently of FruA (Fig. 6A), binding of MrpC was not detected at 4 μM (Fig. 9C, lane 5) or at concentrations ranging up to 20 μM (Fig. S8C, lanes 12-16), and MrpC did not affect MBP-Nla6 DBD or FruA binding (Fig. 9C, lanes 9, 11, 12, 14). Taken together, our results suggest that MrpC regulates *exoA* indirectly both independently of FruA (Fig. 6A) and because MrpC is required for synthesis of FruA (29, 62), which appears to bind cooperatively with Nla6 to the promoter region and increase *exoA-I* transcription.

We discovered that FruA binds to the *exoL-P* promoter region fragment (Fig. 9D, lane 4). Binding of MBP-Nla6 DBD was not detected at 4 μM in the experiment shown in Figure 9D (lane 2), but was detected at 4 μM in two other experiments (Fig. S8D, lanes 2, 8). Interestingly, MBP-Nla6 DBD and FruA appeared to bind cooperatively and the order of addition made no difference (Fig. 9D, lane 6 and Fig. S8D, lanes 4-6), similar to our results for the *exoA-I* fragment. Both fragments also yielded similar results in other experiments; FruA DBD exhibited very little cooperative binding with MBP-Nla6 DBD to the *exoL-P* fragment (Fig. S8D, lane 10) and MrpC did not affect MBP-Nla6 DBD or FruA binding (Fig. 9D, lanes 5, 7). However, the two fragments differed in terms of MrpC binding, which was not detected for the *exoA-I* fragment at MrpC concentrations ranging up to 20 μM (Fig. S8C, lanes 12-16). In contrast, for the *exoL-P* fragment, although binding of MrpC was not detected at 4 μM in the experiment shown in Figure 9D (lane 3), binding was detected at 4 μM and increased at higher concentrations in another experiment (Fig. S8D, lanes 12-16). Therefore, MrpC (as well as FruA and Nla6, which bind cooperatively) appears to directly regulate *exoL-P*, differentiating its regulation from that of *exoA-I* (see Discussion).

We found that MBP-Nla6 DBD, MrpC, and FruA could each bind individually to the *nfsA-H* promoter region fragment (Fig. 9E, lanes 2-7). In combination with MBP-Nla6 DBD at 4 μM, individual binding of MrpC was not detected and the intensity of the shifted complex increased at the lagging edge (Fig. 9E, lane 9), suggestive of cooperative binding. In agreement, MBP-Nla6 DBD at 2 μM appeared to bind cooperatively with MrpC, and the order of addition made little or no difference (Fig. S8E, lanes 4-6). We note that the large sizes of MBP-Nla6 DBD and the DNA fragment likely explain the small difference in migration of the putative MBP-Nla6 DBD/MrpC/DNA complex compared to the MBP-Nla6 DBD/DNA complex (Fig. S8E, lanes 2, 4-6). Like MrpC, FruA appeared to bind cooperatively with MBP-Nla6 DBD (Fig. 9E, lane 10 and Fig. S8E, lanes 10-12). In the mixtures with MBP-Nla6 DBD, individual binding of FruA was detected, suggestive of less cooperativity than in the case of MrpC. FruA DBD did not appear to bind cooperatively MBP-Nla6 DBD (Fig. S8E, lane 16), and FruA did not appear to bind cooperatively with MrpC (Fig. S8E, lanes 20-22). We conclude that MBP-Nla6 DBD, MrpC, and FruA appear to directly regulate *nfsA-H* and we note that the apparent cooperative binding of MBP-Nla6 DBD and MrpC to the *nfsA-H* fragment distinguishes it from the *exoA-I* and *exoL-P* fragments.

Altogether, the results of our EMSAs and transcript measurements suggest that *fadIJ* regulation is similar to that of *dev* (42), *fmg* (31, 43–45), and likely many other genes or operons (41), involving cooperative binding of MrpC and C-signal-activated FruA. In contrast, regulation of the other late-acting operons appears to involve cooperative binding of Nla6 and FruA, and MrpC acting indirectly (*exoA-I*), directly but not binding cooperatively with Nla6 or FruA (*exoL-P*), or binding cooperatively with Nla6 (*nfsA-H*).

### DevI negatively regulates the transcript levels of the late-acting operons

Reporter activity from fusions to *exoC* (61) and *nfsA* (57) was reduced in a *devRS* insertion mutant compared with a WT strain during development. The *exoA* transcript level was also reduced in a *devS* in-frame deletion mutant compared with a WT strain during development (53). To further investigate the effects of mutations in the *dev* operon, which comprises a CRISPR-Cas system (51), on regulation of the late-acting operons, we measured transcript levels in *devI* and *devS* in-frame deletion mutants. Sporulation occurs about 6 h earlier than normal in the *devI* mutant (29, 53), suggesting that DevI delays sporulation of the WT strain. Sporulation is severely impaired in the *devS* mutant (29, 51) due to loss of negative autoregulation of *dev* transcription and resulting overproduction of DevI, which strongly inhibits sporulation (52).

We found that the *exoA*, *exoL*, *nfsA*, and *fadI* transcript levels remain low in the *devS* mutant at 18-30 h PS (Fig. 10). We infer that DevI overproduction in the *devS* mutant inhibits accumulation of transcripts from the late-acting operons. Although Student’s two-tailed *t*-tests rarely yielded *p* < 0.05 comparing transcript levels in the *devS* mutant with the WT strain at the corresponding time PS, this was due to the large variation between biological replicates of the WT strain. We emphasize that transcript levels were low in all biological replicates of the *devS* mutant at all times PS.

**Figure 10.**
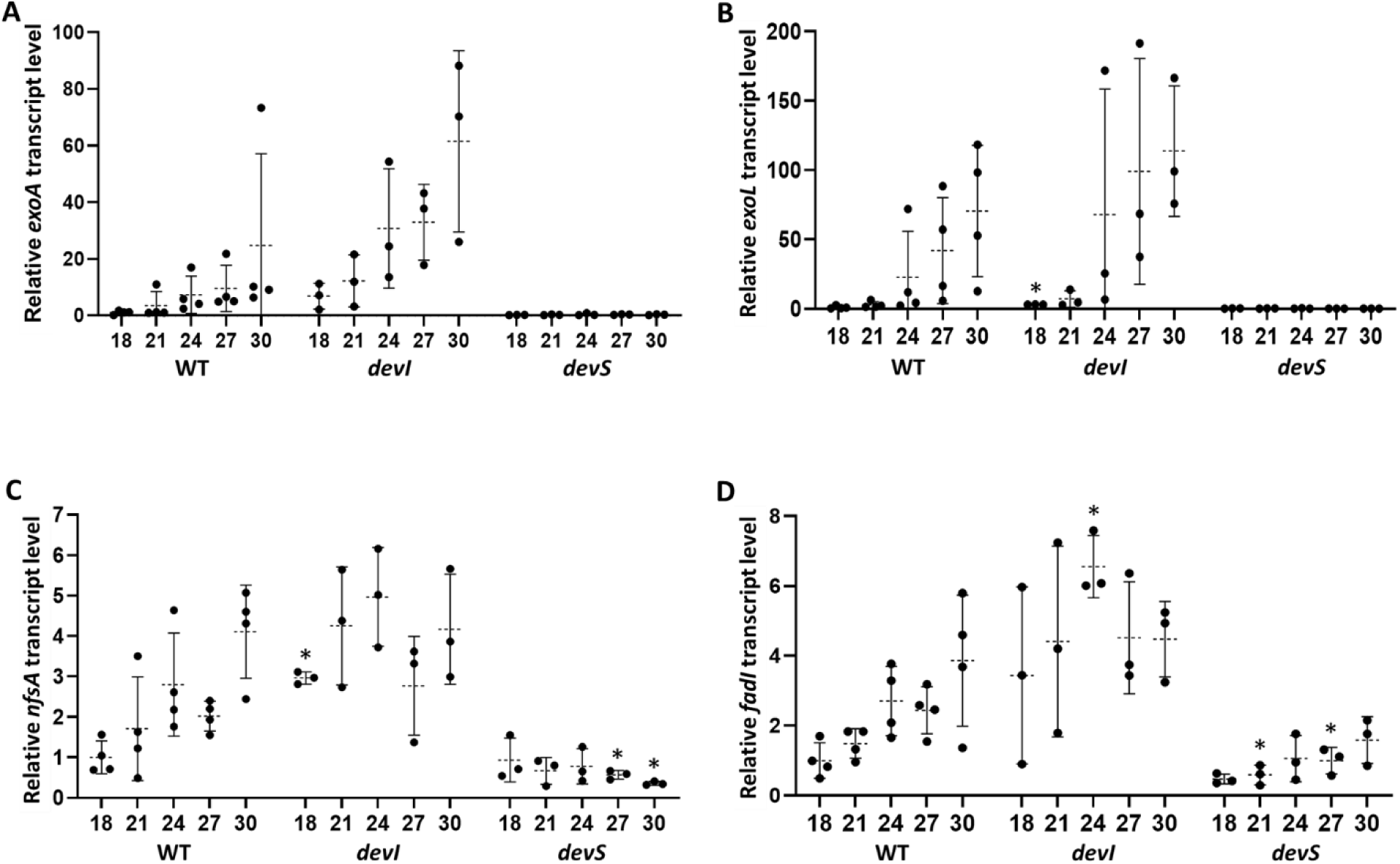
Transcript levels in *devI* and *devS* mutants during *M. xanthus* development. The WT strain and its mutant derivatives were subjected to starvation under submerged culture conditions. Samples were collected at the indicated times PS for measurement of *exoA* (A), *exoL* (B), *nfsA* (C), and *fadI* (D) transcript levels by RT-qPCR. Graphs show the data points and the average of at least three biological replicates, relative to the WT strain at 18 h PS, and error bars indicate one standard deviation. Asterisks indicate a significant difference (*p* < 0.05 in Student’s two-tailed *t*-tests) from the WT strain at the corresponding time PS.

The average transcript levels were greater in the *devI* mutant than the WT strain, except the *nfsA* transcript level was similar in both at 30 h PS (Fig. 10). These results are consistent with the notion that DevI inhibits accumulation of transcripts from the late-acting operons. Here again, Student’s two-tailed *t*-tests rarely yielded *p* < 0.05, but in this case there was large variation between biological replicates of both the *devI* mutant and the WT strain. Even so, taken together, the measurements of transcript levels in the *devI* and *devS* mutants support that DevI is a negative regulator of the late-acting operons.

## Discussion

We investigated the function and regulation of genes whose products act near the end of the *M. xanthus* developmental process to change metabolism of the spore and protect its contents by depositing a polysaccharide coat on its surface. We found that the *exoA-I*, *exoL-P*, and *nfsA-H* operons involved in spore coat biogenesis impact the formation of resistant spores earlier than shown previously, but do not prevent the initial cellular shape change associated with sporulation. We discovered that the regulation of the late-acting operons is unusually complex. It involves three transcription factors that bind to the promoter regions in different combinations, allowing each promoter to be regulated uniquely. Combinatorial regulation of this complexity is common in higher eukaryotes (67–69), but not in bacteria. Our results also show that the late-acting operons are controlled by a CRISPR-Cas system. These systems typically defend bacteria against viral infection, but are often repurposed for a variety of functions, including regulation of endogenous gene expression (70, 71). Collectively, our results suggest that *M. xanthus* evolved complex eukaryotic-like combinatorial control of transcription and a link to a CRISPR-Cas system to thwart viral intrusion while making spores suited to withstand starvation and environmental insults.

### New insights into the functions of the late-acting operons

Our results show that mutations in *exoC*, *nfsA-H*, and *exoL* reduce the formation of sonication-resistant spores beginning at 27 h PS (Fig. 3), but do not prevent the initial cellular shape change associated with starvation-induced sporulation (Fig. S1B). Previously, these mutants were examined for sporulation at 120 h (58, 63, 64). Our findings indicate a much earlier role of the ExoA-I, NfsA-H, and ExoL-P proteins during development than established previously. The *nfsA-H* mutant made about twofold less sonication-resistant spores than the WT strain at 27-48 h, whereas the *exoC* and *exoL* mutants made less than the detection limit (Fig. 3). The milder sporulation defect of the *nfsA-H* mutant compared with the *exoC* and *exoL* mutants (Fig. 3 and Table S1) is consistent with previous reports (58, 63, 64). The *exoC* mutant fails to export spore coat polysaccharide, whereas the *nfsA-H* mutant appears to export it, but fails to assemble it properly, producing an amorphous, unstructured coat that nevertheless appears to provide some heat- and sonication-resistance (58). The *fadI* mutant made about twofold more sonication-resistant spores than the WT strain (Fig. 3). The *fadI* insertion mutant presumably has a reduced rate of fatty acid β-oxidation (55), so perhaps altered metabolism enhanced formation of sonication-resistant spores (Fig. 3), albeit not mature spores at 72 h (Table S1).

Building on the recent observation of TCs in nascent fruiting bodies (46), we devised a method that allowed us to visualize and enumerate TCs in samples of the starved mutants (Fig. S1B). These TCs likely resemble cells of *exo* and *nfs* mutants that fail to complete morphogenesis upon chemical induction of sporulation in liquid culture (58, 59, 64). The lower percentage of TCs we observed for *exoC*, *nfsA-H*, and *exoL* mutants as compared with the WT strain at 24 and 27 h (Fig. S1B) may reflect reduced ability to initiate and/or maintain the cellular shape change associated with sporulation.

### Combinatorial regulation by three signal-responsive transcription factors links expression of late-acting operon to environmental conditions

Eukaryotic-like signaling and gene regulation have been known in *M. xanthus* for decades (72, 73). More recently, combinatorial control, a common theme in eukaryotic GRNs (67–69), emerged from studies of *M. xanthus* gene regulation (7, 35). Combinatorial control allows signal integration by utilizing two or more signal-responsive transcription factors to directly regulate the same gene. Our transcript measurements demonstrate unique regulation of each late-acting operon and our DNA-binding assays implicate combinatorial control of each promoter region by two or three transcription factors (Fig. 11). The late-acting operons are shown twice to indicate differential regulation early (∼6-18 h PS) and late (∼21-30 h) in development (gray boxes at right). The early period approximates when mound formation occurs in the WT strain (Fig. 2, S5, S6, and the cartoon at left in Fig. 11). The late period includes times leading up to and including the beginning of sporulation (Fig. 3 and 11). Our transcript measurements focused on the early (Fig. 7 and 8) or late (Fig. 4, 6, 10, and S2) period and both included 18 h. The results show that transcript levels began to increase markedly by 18 h (*exoA*, *nfsA*, and *fadIJ*) or 24 h (*exoL*) and continued to increase until 30 h. For simplicity, our model ignores this difference in timing of transcript increase and we focus primarily on explaining negative regulation during the early period and positive regulation during the late period. In the model, green arrows and red lines with a barred end indicate positive and negative regulation, respectively, based on our transcript measurements. Solid versus dashed arrows/lines represent our interpretation of our EMSA results as indicative of direct versus indirect regulation. We interpret binding of transcription factors at relatively low concentrations (≤ 4 μM) to a promoter region fragment as indicative of direct regulation, and either no detectable binding or binding at relatively high concentrations (≥ 16 μM) as indicative of indirect regulation. Cartoons below the operons in the model also take into account our DNA sequence analysis of potential binding sites for the transcription factors (Fig. S9), in an effort to illustrate how independent, cooperative, and competitive binding could explain the complex regulation observed. Importantly, much more work will be required to test our model. For example, binding sites will need to be identified and mutated, and the effects on transcription measured.

**Figure 11.**
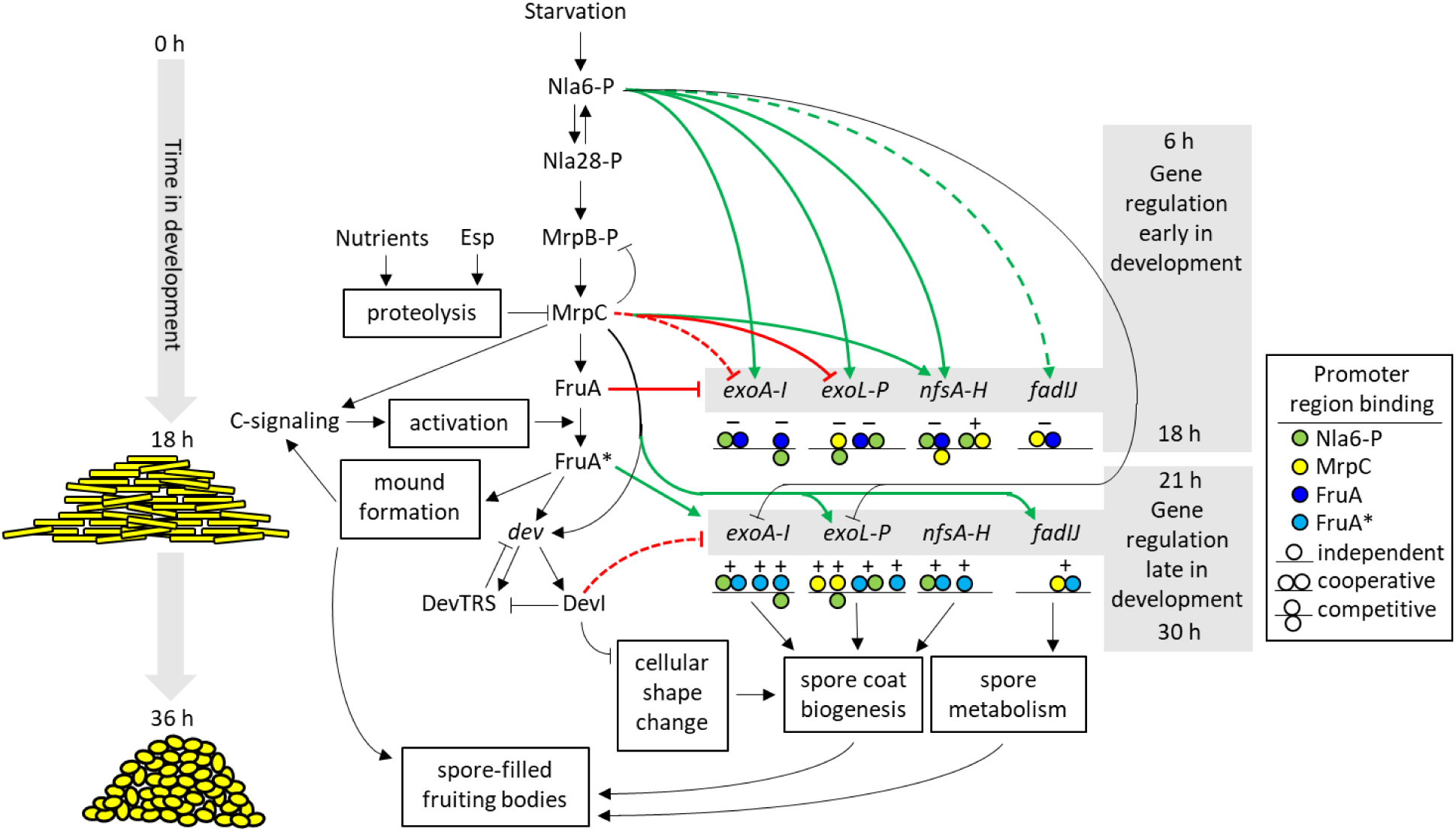
Model for regulation of late-acting operons during *M. xanthus* development. See the Figure 1 legend for explanation of the parts of the GRN that were known prior to this study. The new findings of our study are shown in color and are explained in the text. Briefly, the cartoon at left depicts a timeline of the developmental process with mounds forming by 18 h PS and spore-filled fruiting bodies forming by 36 h. Green arrows indicate positive regulation and red lines with a barred end indicate negative regulation. Solid arrows/lines indicate direct regulation and dashed arrows/lines indicate indirect regulation. Cartoons below the operons illustrate how independent, cooperative, and competitive binding of transcription factors to the promoter regions (see boxed key) could explain negative regulation early in development and positive regulation late in development. These cartoons are further explained in the Supporting Information.

To explain negative regulation of the late-acting operons early in development, our model proposes that unactivated FruA directly opposes stimulation of transcription by Nla6-P at the *exoA-I*, *exoL-P*, and *nfsA-H* promoter regions (Fig. 11). This model could explain a large increase in the *exoA-I* and *exoL-P* transcript levels between 0 and 1 h PS followed by a comparable decrease by 12 h in a WT strain (63). FruA is present at 6 h and its level increases threefold by 12 h (Fig. S4A). FruA* appears to be present by 18 h (29), but the kinetics of FruA activation are unknown. Presumably, mound formation brings cells into proximity, enabling C-signal-dependent activation of FruA (19, 46), so FruA* increases and unactivated FruA decreases during mound formation, which occurs primarily between 12 and 18 h under our conditions (Fig. S5). Therefore, we expect unactivated FruA to be abundant by 6 h (possibly earlier), perhaps opposing Nla6-P stimulation after the initial increase of *exoA-I* and *exoL-P* transcript levels (63).

Our model proposing negative regulation by unactivated FruA early in development (Fig. 11) is consistent with two prior reports of negative regulation by FruA. Expression of a reporter fused to the *nfsA-H* promoter region was elevated in a *fruA* mutant compared with a WT strain at 12-18 h PS (57). While our work was in progress, transcriptomic analysis identified > 1500 genes with an elevated transcript level in the *fruA* mutant compared to the WT strain at 12 h (6). Based on the data of McLoon *et al.* (6), the transcript levels of all genes in the late-acting operons were elevated in the *fruA* mutant relative to the WT strain at 12 h (Table S2). In our study, the *exoA-I*, *exoL-P*, and *nfsA-H* transcript levels were elevated in the *fruA* mutant relative to the WT strain at 18 h (Fig. 4, 6, and 7), but not at 12 h (Fig. 7), suggesting a temporal difference in development between the studies. The *fadI* transcript level was lower in the *fruA* mutant than in the WT strain at all times tested in our experiments (Fig. 4D, 6D, and 7D), but its level in the *fruA* mutant exceeded that in the *csgA* mutant by a small, reproducible amount at 18 h (Fig. 4D, *p* = 0.002 in a Student’s two-tailed *t*-test), supporting weak negative regulation by unactivated FruA (Fig. 11). Our model proposes that unactivated FruA directly opposes stimulation of *fadIJ* transcription by cooperative binding of MrpC and FruA* early in development (Fig. 11 cartoon), rather than opposing stimulation by Nla6-P, for two reasons. First, a high concentration of MBP-Nla6 DBD was required to detect binding to the *fadIJ* promoter region fragment (Fig. S8B), suggesting indirect regulation by Nla6-P (Fig. 11). Second, our analysis revealed potential binding sites for MrpC and FruA or FruA*, but not for Nla6, upstream of the *fadIJ* promoter (Fig. S9A).

Our model also proposes that MrpC affects transcription of the *exoA-I*, *exoL-P*, and *nfsA-H* operons early in development (Fig. 11). The *exoA* transcript level was on average at least twofold greater in the *mrpC* mutant than the *fruA* mutant at 18 h PS (Fig. 6A and 7A), suggesting negative regulation by MrpC independently of FruA, but we did not detect MrpC binding to the *exoA-I* promoter region fragment (Fig. S8C) or potential binding sites (Fig. S9D). Therefore, we propose that MrpC indirectly decreases transcription of the *exoA-I* operon early in development of the WT strain (Fig. 11). In contrast, our results imply that MrpC directly decreases *exoL-P* transcription and directly increases that of *nfsA-H* early in development (Fig. 6, 7, 9, S8, and 11).

The cartoons below the *exoA-I*, *exoL-P*, and *nfsA-H* operons in Figure 11 propose that unactivated FruA and MrpC engage in cooperative or competitive binding with Nla6-P or each other primarily to inhibit transcription early in development of the WT strain. Our EMSAs (Fig. 9 and S8) and our analysis revealing adjacent potential binding sites (Fig. S9) support the cooperative binding depicted. We infer competitive binding at overlapping potential binding sites (Fig. S9). The arrangement of transcription factors in the cartoons (Fig. 11) mimics the arrangement of potential binding sites (Fig. S9) proposed to play important roles (Supporting Information, additional discussion). Not all the potential binding sites shown in Figure S9 are depicted in the Figure 11 cartoons.

To explain positive regulation of the late-acting operons late in development, our model proposes that C-signaling switches FruA from a negative regulator to a positive regulator, FruA* (Fig. 11). The switch is most evident comparing the *nfsA* transcript levels in the *fruA* mutant with those in the WT strain as development proceeds (i.e., the transcript level increases in the WT strain and decreases in the *fruA* mutant) (Fig. 4C). The switch is less evident from a comparison of *exoA* transcript levels due to large variation between biological replicates of the WT strain late in development, but on average the transcript level increases in the WT strain and decreases in the *fruA* mutant (Fig. 4A). Our findings that ectopic FruA expression in the *mrpC* mutant allowed formation of immature mounds (Fig. S5) and greatly increased the *exoA* transcript level (Fig. 7A) also support a switch to stimulation of transcription by FruA* (Fig. 11). Similar findings suggest that FruA* increases the *exoL* transcript level (Fig. 7B and 11). However, the *exoL* transcript level does not decrease in the *fruA* mutant late in development (Fig. 4B). Therefore, we propose that MrpC can act as a direct positive regulator independently of FruA* (Fig. 11). In contrast, our results suggest that cooperative binding of MrpC and FruA* stimulates *fadIJ* transcription late in development (Fig. 11).

The cartoons in Figure 11 shown below the operons late in development reflect our EMSA results (Fig. 9 and S8) and our analysis of potential binding sites (Fig. S9), using similar reasoning as explained above, except we propose that FruA* may differ from unactivated FruA in terms of affinity for potential binding sites and the effect of cooperative binding with Nla6-P or dephosphorylated Nla6 (Supporting Information, additional discussion). In short, we propose that MrpC and FruA* bind cooperatively near the *fadIJ* promoter to stimulate transcription late in development, whereas binding of MrpC and unactivated FruA farther upstream weakly inhibits transcription stimulation earlier (Fig. 11 cartoons). The cartoons below the *exoA-I*, *exoL-P*, and *nfsA-H* operons propose that FruA* and MrpC bind independently and/or engage in cooperative or competitive binding with Nla6-P (or dephosphorylated Nla6, which is not shown for simplicity) primarily to stimulate transcription late in development of the WT strain.

We conclude that each late-acting operon is uniquely regulated by two or three transcription factors acting combinatorially. The positions and affinities of binding sites for Nla6-P or dephosphorylated Nla6, MrpC, and FruA or activated FruA* in each promoter region determine the transcriptional output from independent, cooperative, and competitive binding. This regulatory complexity was masked in transcriptomic studies. The four late-acting operons we investigated were clustered in one (5) or two (4, 6) groups with hundreds of other genes, but our transcript measurements in several mutants revealed unique regulation of each operon (Fig. 4, 6, and 7), our DNA-binding assays showed different binding of transcription factors to each promoter region (Fig. 9 and S8), and our analysis of potential binding sites uncovered diverse, complex architectures (Fig. S9). Collectively, our results and the transcriptomic studies predict that tremendous regulatory complexity will be uncovered by genome-wide measurements of transcript levels in mutants, transcription factor binding studies, and bioinformatic analyses.

Since combinatorial control allows signal integration, what are the signals to which the late-acting operons respond? Starvation triggers a cascade of EBPs that includes Nla6-P and Nla28-P (32), and leads to the production of MrpB-P (74), another EBP that in turn leads to production of MrpC (75) (Fig. 11). However, more specific molecular signals controlling these steps are unknown. In particular, it is unknown whether Nla6 responds to different signals early and late in development, or how it switches from positive to negative regulation of the *exo* operons (63) (Fig. 11). Nutrients and an unknown signal sensed by the Esp system stimulate proteolysis of MrpC (Fig. 11) to delay or halt development (36–39). The presumed advantage of such a developmental checkpoint is to ensure that cellular physiology and interactions with other cells, as well as environmental conditions, warrant commitment to spore formation (38, 76, 77). C-signaling appears to require the movement of cells into alignment or possibly into contact, and lead to the formation of active FruA* (18–20, 28, 29, 46). Early in development, unactivated FruA appears to function as a global negative regulator (6) (Fig. 11), preventing premature production of some proteins that would be detrimental during mound formation (e.g., Nfs and Exo proteins involved in polysaccharide export), in addition to conserving cellular resources. Altogether, our knowledge about the specific signals to which the late-acting operons respond is quite limited, but in general, nutrient limitation and movement of cells into proximity appear to play important roles, and are likely influenced by many other factors in the environment of the developing population.

We propose that combinatorial control of the late-acting operons and hundreds of other genes by signal-responsive transcription factors allows proper temporal and spatial expression of genes involved in spore metabolism, coat biogenesis, and other functions in order to produce spores that are densely packed in fruiting bodies and suited to withstand environmental insults. Numerous studies show that environmental conditions affect the resistance and surface characteristics of endospores from *Bacillus* species (78–80). In *Bacillus subtilis*, several transcription factors positively and negatively fine-tune the expression of hundreds of genes during sporulation (81–83), ensuring that the resulting spores are endowed with resistance and surface properties tailored for their environment (84–86). Understanding how ecology and evolution impact the developmental GRN of *M. xanthus* and related species is an ongoing and future challenge (6, 87, 88).

### The DevI component of a CRISPR-Cas system is a negative regulator of the late-acting operons

Our results confirm and extend the role of DevI as a negative regulator of late-acting operons during development. We confirmed that the *exoA* transcript level is very low in the *devS* null mutant (53) and we observed similar results for the *exoL*, *nfsA*, and *fadI* transcript levels (Fig. 10). The cumulative effect of failure to express the late-acting operons may explain the inability of the *devS* mutant to undergo normal cellular shape change (53). Alternatively, one or more other genes subject to negative regulation by DevI overproduction may be responsible. We also found that transcript levels from the late-acting operons were slightly greater on average in a *devI* mutant than in the WT strain, supporting our conclusion that DevI is a negative regulator of the late-acting operons (Fig. 11); however, the mechanism of negative regulation by DevI is unknown.

The *dev* CRISPR-Cas system appears to have evolved recently (52), perhaps in niches where protection from bacteriophage infection and/or delayed sporulation were advantageous (51, 53). Most natural isolates of *M. xanthus* lack *devI* and their *dev* promoter region is predicted to be nonfunctional (52). The *dev* operon is dispensable for development of the WT laboratory strain used in this work (53). However, the *dev* CRISPR-Cas system may protect against intrusion by bacteriophage Mx8 during development, since a 37-bp sequence in the CRISPR matches a sequence in the Mx8 integrase gene (51). This hypothesis remains to be tested, as do the potential benefits of such protection.

In summary, our investigation uncovered complex regulation of genes involved in spore metabolism and spore coat biogenesis by three transcription factors and a CRISPR-Cas component during *M. xanthus* development. Further elucidating the signals to which Nla6, MrpC, and FruA respond, understanding how these transcription factors interact at promoter regions, and elucidating how DevI inhibits the cellular shape change associated with sporulation are important goals for the future.

## Experimental procedures

### Bacterial strains, plasmids and primers

The strains, plasmids and primers used in this study are listed in Table S3.

*M. xanthus* strain MSS2 with P*_van_-fruA* integrated ectopically was constructed by electroporating (89) pSS10 into strain DK1622, selecting transformants on CTT agar (1% Casitone, 10 mM Tris-HCl [pH 8.0], 1 mM KH_2_PO_4_-K_2_HPO_4_, 8 mM MgSO_4_ [final pH 7.6] solidified with 1.5% agar) containing 15 μg/mL tetracycline (90), and verification by colony PCR using primers pMR3691 MCS G-F and pMR3691 MCS G-R.

*M. xanthus* strain MSS10 with a plasmid insertion mutation in *nla6* was constructed by electroporating pSS11 into strain DK1622, selecting transformants on CTT agar containing tetracycline (15 µg/mL), and verification by colony PCR using primers PMR3487 Rev, Nla6 Fwd4, and Nla6 Fwd5. To construct pSS11, primers Nla6 Fwd and Nla6 Rev were used to generate a PCR product using chromosomal DNA from *M. xanthus* strain DK1622 as a template. The product was combined with DNA amplified from pMR3487 using primers PMR3487G Fwd and PMR3487G Rev primers, and a Gibson assembly reaction was used to enzymatically join the overlapping DNA fragments (91). The reaction mixture was transformed into *E. coli* strain DH5α, transformants were selected at 37°C on Luria-Bertani (LB) (92) 1.5% agar containing 15 μg/mL tetracycline, and the cloned DNA sequence was verified using primers 3487 seq Fwd1, 3487 seq Fwd2, 3487 seq Fwd3, 3487 seq Fwd4, and 3487 seq Fwd5.

To provide a DNA template for amplification of the *nfsA-H* promoter region, pSS14 was constructed as follows. Primers Nfs −290G and Nfs +83G were used to generate a PCR product using chromosomal DNA from *M. xanthus* strain DK1622 as a template. The product was combined with DNA amplified from pMR3487, the overlapping DNA fragments were joined using a Gibson assembly reaction, the reaction mixture was transformed into *E. coli* strain DH5α, transformants were selected, and the cloned DNA sequence was verified as described above for pSS11.

To express MBP-Nla6 DBD for purification, primers Nla DBD For and Nla DBD Rev were used to generate a PCR product using *nla6* DNA as a template. The PCR product was digest with *Sbf*I and ligated to *Xmn*I-*Sbf*I-digested pMAL-c5x (NEB), resulting in pMAL-c5x/MBP-ΔCNla6 DBD, which lacked a CG bp at the beginning of the Nla6 DBD-encoding segment. The missing CG bp was added by performing site-directed mutagenesis using the QuikChange strategy (Stratagene) with primers MBP-Nla6 add C fwd and MBP-Nla6 add C rev, resulting in pMAL-c5x/MBP-Nla6 DBD, and the DNA sequence was verified using primer mbp fwd colpcr.

### Growth and development of M. xanthus

Strains of *M. xanthus* were grown at 32°C in CTTYE liquid medium (CTT medium with 0.2% yeast extract) with shaking at 350 rpm. CTT agar was used for growth on solid medium and was supplemented with 40 µg/mL of kanamycin sulfate or 15 µg/mL of tetracycline as required. Fruiting body development under submerged culture conditions was performed using MC7 (10 mM morpholinepropanesulfonic acid [MOPS; pH 7.0], 1 mM CaCl_2_) as the starvation buffer as described previously (38). Briefly, cells in the mid-exponential phase of growth were collected by centrifugation, the supernatant was removed, and the cells were resuspended in MC7 buffer at a density of approximately 1,000 Klett units. A 96-μL sample (designated *T*_0_) was removed and 4 µL of glutaraldehyde was added from a 50% stock solution to achieve a 2% final concentration. The sample was stored at 4°C for at least 24 h in order to fix the cells for later quantification of total cells as described below. For each developmental sample, 1.5 mL of the 1,000-Klett-unit cell suspension plus 10.5 mL of MC7 buffer was added to an 8.5-cm-diameter plastic petri plate. Upon incubation at 32°C, cells adhere to the bottom of the plate and undergo development. At the indicated times, developing populations were photographed using a Leica Wild M8 microscope equipped with an Olympus E-620 digital camera.

### Sample collection

At the indicated times PS, the MC7 buffer overlay was replaced with 5 mL of fresh MC7 buffer or 5 mL of MC7 buffer containing 50 μg/mL of rifampicin to inhibit transcription for measurement of transcript degradation rates. Developing cells were scraped from the bottom of the plates, the entire contents were collected in a 15-mL centrifuge tube, and samples were mixed thoroughly as described previously (29). A 96-μL sample was removed, glutaraldehyde was added, and the sample was stored as described above for the *T*_0_ sample, fixing the cells for quantification of total cells as described below. A 400 µL sample was removed and stored at - 20°C for measurement of sonication-resistant spores as described below. For the experiments shown in Figures 8 and S4, a 100 μl sample was removed and added to an equal volume of 2× sample buffer (0.125 M Tris-HCl [pH 6.8], 20% glycerol, 4% sodium dodecyl sulfate [SDS], 0.2% bromophenol blue, 0.2 M dithiothreitol), boiled for 5 min, and stored at −20°C for immunoblot analysis. Immediately after collecting the samples just described, the rest of the 1,000-Klett- unit cell suspension was mixed with 0.5 mL of RNase stop solution (5% phenol [pH < 7] in ethanol), followed by rapid cooling in liquid nitrogen until almost frozen, centrifugation at 8,700 × *g* for 10 min at 4°C, removal of the supernatant, freezing of the cell pellet in liquid nitrogen, and storage at −80°C until RNA extraction.

### Quantification of total cells, transitioning cells, and sonication-resistant spores

The total number of cells, including rods, elliptical TCs, and round spores, was determined using the glutaraldehyde-fixed samples collected as described above. Each sample was thawed and mixed by vortexing and pipetting, diluted with MC7 buffer, sonicated for 10 s, and then all cells were counted microscopically as described previously (29), except noting the number of cells that were elliptical or round. The number of sonication-resistant spores in the 400-µL samples collected as described above was quantified as described previously (38). These samples were not glutaraldehyde-fixed and were sonicated for 10 s intervals three times with cooling on ice in between, leaving only round spores (i.e., neither rods nor elliptical TCs were observed). The total cell number minus the number of sonication-resistant spores was designated the number of sonication-sensitive cells (consisting primarily of rods and a small percentage of TCs) and was expressed as a percentage of the total cell number in the corresponding *T*_0_ sample (consisting only of rods). The number of cells that were elliptical or round in the glutaraldehyde-fixed sample minus the number of sonication-resistant spores in the corresponding sample that was not glutaraldehyde-fixed, was designated the number of TCs and was also expressed as a percentage of the total cell number in the corresponding *T*_0_ sample.

### RNA extraction and analysis

RNA was extracted using the hot-phenol method, followed by digestion with DNase I (Roche) as described previously (36). The RNA (1 µg) was then subjected to cDNA synthesis using Superscript III reverse transcriptase (InVitrogen) and random primers (Promega), as instructed by the manufacturers. In parallel, RNA (1 µg) was subjected to cDNA synthesis reaction conditions without Superscript III reverse transcriptase, as a control. One µl of cDNA at the appropriate dilution (as determined empirically) and 20 pmol of each primer were subjected to qPCR in a 25 µl reaction using 2× reaction buffer as described previously (29). qPCR was done in quadruplicate for each cDNA using a LightCycler® 480 System (Roche). In parallel, a standard curve was generated for each pair of qPCR primers using genomic DNA (gDNA) from *M. xanthus* WT strain DK1622 and gene expression was quantified using the relative standard curve method (user bulletin 2; Applied Biosystems). 16S rRNA was used as the internal standard for each sample. The primers used for RT-qPCR analysis of 16S rRNA (93) and the late-acting operons (39) have been described previously.

Relative transcript levels for mutants are the average of three biological replicates after each replicate was normalized to the transcript level observed for one replicate of the WT strain at the earliest time point (i.e., either 6 or 18 h PS) in the same experiment (29). Transcript levels for the WT strain at other times PS were likewise normalized to that observed for the WT strain at the earliest time point in the same experiment. For the WT strain at the earliest time point, the transcript levels of at least three biological replicates from different experiments were normalized to their average, which was set as 1.

To measure transcript degradation rates, samples were collected immediately after addition of rifampicin (designated t_0_) and 8 and 16 min later for RNA extraction and analysis as described above. For each biological replicate the transcript levels after 8 and 16 min were normalized to the transcript level at t_0_, which was set as 1, and the natural log of the resulting values was plotted versus minutes after rifampicin treatment (29). The slope of a linear fit of the data was used to compute the mRNA half-life.

### Expression and purification of proteins

To express His-tagged proteins, *E. coli* strain BL21(DE3) was transformed with pET11a/FruA- DBD-H_8_, pET11km/FruA-H_6_, or pPH158 (for H_6_-MrpC) and transformants were selected on LB 1.5% agar containing 50 μg/mL ampicillin or kanamycin sulfate. In each case, a single colony was used to inoculate 10 mL of LB supplemented with antibiotic, followed by overnight incubation at 37°C with shaking. Five mL of overnight culture was used to inoculate 500 mL of the same medium, followed by continued shaking at 37°C until the culture reached 60-80 Klett units. IPTG (1 mM final concentration) was added to induce synthesis of the recombinant protein. After 2 h, cells were harvested as reported previously (94) and stored at −80°C.

To purify His-tagged proteins, a modification of a protocol described previously (95) was used. Each cell pellet was resuspended in 35 mL of lysis buffer (50 mM Na-phosphate buffer pH 8.0, 500 mM NaCl, 14 mM β-mercaptoethanol, protease inhibitors [1 Roche Mini EDTA-free tablet/10 mL]). Cells were disrupted by sonication (4 times for 1 min with intermittent cooling on ice). After centrifugation at 18,000 × *g* for 10 min at 4°C, the supernatant was mixed with 5 mL of lysis buffer containing 10% w/v Triton X-100, 1 mL of 1 M imidazole (pH 8.0), enough lysis buffer for 50 mL total, and finally 0.5 mL of Ni-NTA agarose (Qiagen) that had been washed 3 times with lysis buffer containing 20 mM imidazole. The mixture was rotated for 1 h at 4°C. The Ni-NTA agarose was collected by centrifugation at 700 × *g* for 3 min at 4°C and washed 4 times with 50 mL of lysis buffer containing 20 mM imidazole, 0.5% w/v Triton X-100, and 25% w/v glycerol. The Ni-NTA agarose was rotated for 30 min at 4°C with 10 mL of elution buffer (50 mM Na-phosphate buffer pH 8.0, 500 mM NaCl, 1.4 mM β-mercaptoethanol, protease inhibitors [1 Roche Mini EDTA-free tablet], 250 mM imidazole, 0.5% w/v Triton X-100, 25% w/v glycerol). Eluates were dialyzed twice at 4°C against 1 L of storage buffer (10 mM Tris-HCL pH 8.0, 100 mM NaCl, 1 mM β-mercaptoethanol, 10% w/v glycerol) and aliquots were stored at - 80°C. The protein concentration was determined using a Bradford (96) assay kit (Bio-Rad).

To express MBP-Nla6 DBD, *E. coli* strain BL21(DE3) was transformed with pMAL-c5x/MBP- Nla6 DBD and transformants were selected on LB 1.5% agar containing 100 μg/mL ampicillin. Five colonies were used to inoculate 80 mL of LB supplemented with antibiotic and 0.2% dextrose, followed by incubation at 37°C with shaking at 350 rpm until the culture reached 60 Klett units. IPTG (0.3 mM final concentration) was added to induce synthesis of the recombinant protein. After 2 h, cells were harvested, the cell pellet was resuspended in 10 mL of column buffer (20 mM Tris-HCl pH 7.5, 200 mM NaCl, 1 mM EDTA, protease inhibitors [1 Roche Mini EDTA-free tablet], 10 mM β-mercaptoethanol), and the cell suspension was stored at −20°C.

MBP-Nla6 DBD was purified according to instructions in the pMAL^TM^ Protein Fusion & Purification System manual (NEB). Briefly, the cell pellet was thawed and cells were disrupted by sonication (8 times for 15 s with intermittent cooling on ice). After centrifugation at 18,000 × *g* for 20 min at 4°C, the supernatant was loaded onto a 1 mL amylose gravity-flow column, which had been prewashed with 5 mL of column buffer. The loaded column was washed with 12 mL of column buffer, then eluted with 2.5 mL of column buffer supplemented with 10 mM maltose, collecting fractions of 4 drops each (approximately 200 μL). Fractions were subjected to SDS-PAGE with Coomassie Blue staining. Fractions 4-9, with maximal purified MBP-Nla6 DBD, were pooled, dialyzed as described above, and aliquots were stored at −80°C. The protein concentration was determined by measuring the absorbance at 280 nm and using the calculated extinction coefficient of MBP-Nla6 DBD (1.32 [mg/mL]^-1^cm^-1^).

### Electrophoretic mobility shift assays

^32^P-labelled DNA fragments were generated by PCR using primers labelled with [γ-^32^P]ATP using T4 polynucleotide kinase (NEB) according to the manufacturer’s instructions. The templates for PCR were either plasmid DNA or gDNA from *M. xanthus* WT strain DK1622. The primers and templates were as follows: primers LK1298 and LK1331 with template pPV391 for *dev*, primers *exoA-I* −171 fwd and *exoA-I* −1 rev with template gDNA for *exoA-I*, primers *fadIJ* −214 fwd and *fadIJ* −5 rev with template gDNA for *fadIJ*, primers *exoL-P* −217 fwd and *exoL-P* +56 rev with template gDNA for *exoL-P*, and primers *nfsA-H* −166 fwd and *nfsA-H* +35 with template pSS14 for *nfsA-H.* The labeled DNA fragments were purified by electrophoresis on 15% polyacrylamide gels, followed by visualization using autoradiography, excision, and overnight elution by soaking in TE buffer (10 mM Tris-HCl pH 8.0, 1 mM EDTA) as described previously (94).

Binding reactions (10 μL) were performed as reported previously (94), except the reaction mixtures were incubated at 37°C for 10 min prior to loading on 8% polyacrylamide gels. To determine whether the order of protein addition was important, one protein was added to the reaction mixture during an initial 10 min incubation at 37°C, followed by addition of a second protein and another 10 min incubation at 37°C, prior to loading on 8% polyacrylamide gels. Gels were dried and exposed to X-ray film for autoradiography as described (94).

## Acknowledgements

We thank Monique Floer for advice about high-throughput qPCR and for use of the LightCycler® 480 System. We thank Anthony Garza, Lawrence Shimkets, Ramya Rajagopalan, Wenyuan Shi, Sumiko Inouye, Montserrat Elias-Arnanz, and Penelope Higgs for sharing strains. We thank Muqing Ma and Anthony Garza for constructing pMAL-c5x/MBP-ΔCNla6 DBD. This work was supported by the National Science Foundation (awards MCB-1411272 and IOS-1951025) and by salary support for L.K. from Michigan State University AgBioResearch.

## Author contributions

Conception or design of the study: LK, SS

Acquisition of the data: SS, LK

Analysis or interpretation of the data: SS, LK

Writing of the manuscript: LK, SS

