## supporting information for "Regulation of late-acting operons by three transcription factors and a CRISPR-Cas component during *Myxococcus xanthus* development"

### Discussion

#### *Cartoons depicting promoter region binding by transcription factors*

We identified potential transcription factor binding sites (see Experimental procedures below), which are shown in Figure S9. Based on the potential binding sites, the results of our DNA-binding assays (Fig. 9 and S8), and the results of our transcript measurements in mutants and the WT strain (summarized in Fig. 11), we created cartoons shown in Figure 11 to illustrate how promoter binding by transcription factors could explain regulation of the late-acting operons. The cartoons are explained in the Discussion main text and the legend of Figure 11. Below, we discuss additional information that supports the cartoons.

The cartoon below the *fadIJ* operon early in development depicts negative regulation due to cooperative binding of unactivated FruA and MrpC (Fig. 11). We identified potential FruA binding sites centered at -70.5 and -121.5 adjacent to a potential MrpC binding site centered at -97 (Fig. S9A). We speculate that cooperative binding of unactivated FruA and MrpC to these sites is nonproductive for *fadIJ* transcription and inhibits cooperative binding of MrpC and FruA* to sites centered at -63.5 (i.e., TGACGCGGCGCATC, which is not identified in Fig. S9A due to three mismatches with the consensus sequence and therefore is predicted to be a low-affinity site) and -45.5, respectively, which can stimulate *fadIJ* transcription late in development (Fig. 11). This hypothesis is attractive since cooperative binding of MrpC and FruA* at similar positions appears to stimulate *fmgBC* (1), *fmgD* (2), and *fmgE* (3) transcription late in development. Moreover, the proposed negative regulation of *fadIJ* is analogous to several examples. Negative regulation of *fmgE* appears to involve alternative cooperative binding of FruA* and MrpC to a higher affinity upstream site versus to a lower affinity promoter-proximal site necessary for transcription (3). Negative regulation of *fmgD* (2) and *mrpC* (4) appears to involve competitive binding of MrpC to overlapping sites for FruA* and MrpB-P binding, respectively, which would otherwise lead to increased transcription. For simplicity, the cartoons in Figure 11 show only cooperative binding of MrpC and unactivated FruA early in development, and cooperative binding of MrpC and FruA* at different positions later.

The cartoon below the *nfsA-H* operon early in development depicts negative regulation due to unactivated FruA binding cooperatively with Nla6-P and competitively with MrpC to inhibit potential stimulation of transcription due to cooperative binding of Nla6-P and MrpC (Fig. 11). The best match to the consensus sequence for MrpC binding is centered at -29 (Fig. S9B), consistent with the location of an MrpC-binding site identified by ChIP-seq analysis (5) (Fig. S7D). A sequence matching the Nla6 half-site consensus sequence (6) is centered at -42.5 (Fig. S9B) and appears to be well-positioned for cooperative binding with MrpC (Fig. 9E and S8E) that could increase transcription. However, MBP-Nla6 DBD also binds cooperatively with FruA to the promoter region (Fig. 9E and S8E). The best match to the consensus sequence for FruA binding is centered at -82.5, but two nearby sequences have just one mismatch, and any of the three might explain cooperative binding with MBP-Nla6 DBD owing to a potential half-site centered at -100.5 (Fig. S9B). In this case, cooperative binding involving unactivated FruA and Nla6-P could explain negative regulation (Fig. 11 cartoon) by sequestering Nla6-P and thus inhibiting cooperative binding of MrpC and Nla6-P that would otherwise lead to increased transcription (e.g., as in the *fruA* mutant at 18 h PS; Fig. 6C and 7C). The three potential FruA binding sites overlap with four potential MrpC binding sites (Fig. S9B), so competitive binding of unactivated FruA and MrpC could also play a role in negative regulation by hindering recruitment of MrpC to the promoter region early in development (Fig. 11 cartoon). The cartoon below the *nfsA-H* operon late in development depicts positive regulation due to possible cooperative binding of FruA* and Nla6-P (presumably at the same location) that could stimulate (rather than inhibit) transcription due to a functional change in FruA* relative to unactivated FruA, and we also include the possibility that independent binding of FruA* (e.g., due to a change in its interaction with Nla6-P or dephosphorylated Nla6) stimulates transcription (Fig. 11).

The cartoon below the *exoL-P* operon early in development depicts negative regulation due to competitive binding of MrpC and Nla6-P because potential binding sites centered at -99 and -89.5, respectively, partially overlap (Fig. S9C), so MrpC may interfere with transcriptional activation by Nla6-P (Fig. 11). It also shows cooperative binding of unactivated FruA and Nla6-P since two sets of potential binding sites (Fig. S9C, centered at -130 and -62 for FruA, and at -115.5 and -46.5 for Nla6) are well-positioned to explain the cooperative binding we observed (Fig. 9D and S8D), which may inhibit transcriptional activation by Nla6-P (Fig. 11). The cartoon below the *exoL-P* operon late in development depicts positive regulation due to possible cooperative binding of FruA* and Nla6-P, or independent binding of FruA*, that could stimulate transcription (Fig. 11) as proposed above for the *nfsA-H* operon. The cartoon also shows possible competitive binding of MrpC and Nla6-P that may explain positive regulation by MrpC due to inhibition of negative regulation by Nla6-P (or dephosphorylated Nla6), and we include the possibility that independent binding of MrpC stimulates transcription (Fig. 11).

The cartoon below the *exoA-I* operon early in development depicts negative regulation due to competitive binding of unactivated FruA and Nla6-P since several potential binding sites overlap (Fig. S9D, centered at -42 and -83 for FruA, and at -31.5, -43.5, and -88.5 for Nla6), so unactivated FruA may interfere with transcriptional activation by Nla6-P (Fig. 11). It also shows cooperative binding of unactivated FruA and Nla6-P because potential binding sites centered at -71 and -88.5, respectively, are well-positioned to explain the cooperative binding (Fig. S9D) we observed (Fig. 9C and S8C), which may inhibit transcriptional activation by Nla6-P (Fig. 11). The cartoon below the *exoA-I* operon late in development depicts positive regulation due to possible cooperative binding of FruA* and Nla6-P, or independent binding of FruA*, that could stimulate transcription (Fig. 11) as proposed above for the *nfsA-H* and *exoL-P* operons. The cartoon also shows possible competitive binding of FruA* and Nla6-P that may explain positive regulation by FruA* due to inhibition of negative regulation by Nla6-P (or dephosphorylated Nla6), and we include the possibility that independent binding of MrpC stimulates transcription (Fig. 11).

### Experimental procedures

#### *Immunoblot analysis*

Samples were subjected to a semi-quantitative method of immunoblot analysis as described previously (7). Briefly, equal volumes (10 μL for measurement of MrpC and 15 μL for measurement of FruA) of samples were subjected to SDS-PAGE and immunoblotting along with a sample of *M. xanthus* WT strain DK1622 collected at 6 h PS as an internal control for normalization of signal intensities across immunoblots. Unsaturated signals were detected, quantified, and normalized to the internal control. Each signal was divided by the total protein concentration of a corresponding sample that had been sonicated for measurement of sonication-resistant spores. After removal of the sample for spore quantification, the remaining portion was centrifuged (10,000 × *g* for 1 min) to pellet cell debris and the total protein concentration of the supernatant was determined using a Bradford (8) assay kit (Bio-Rad). The resulting values of normalized signal intensity/total protein concentration were further normalized to the average value for all biological replicates of WT at 6 h PS, which was set as 1.

***Identification of potential transcription factor binding sites***

The DNA sequences of promoter region fragments used in EMSAs were searched for matches to transcription factor binding site consensus sequences using the DNA Pattern Find program (https://www.bioinformatics.org/sms2/dna_pattern.html) of the Sequence Manipulation Suite (9).

**Table S1 Cell and spore numbers**

| Strain | Sonication-sensitive cells at $T_0$ ($10^7$/mL) | Sonication-resistant spores at $T_{48}$ ($10^7$/mL) | Mature spores at $T_{72}$ ($10^6$/mL) |
|---|---|---|---|
| WT | 140 ± 7 | 2 ± 1 | 2.0 ± 0.1 |
| *exoC* | 140 ± 9 | < 0.05 | < 0.00001 |
| *nfsA-H* | 140 ± 7 | 1.4 ± 0.1 | 0.3 ± 0.1 |
| *exoL* | 160 ± 14 | < 0.05 | < 0.00001 |
| *fadI* | 140 ± 6 | 4.3 ± 0.2 | 2 ± 1 |
| *mrpC* $P_{van}$-fruA | 120 ± 22 | < 0.05 | < 0.00001 |
| $Km^r$ *nla6* | 120 ± 15 | < 0.05 | < 0.00001 |
| $Tc^r$ *nla6* | 110 ± 16 | < 0.05 | < 0.00001 |

The WT strain and its indicated mutant derivatives were subjected to starvation under submerged culture conditions. Rod-shaped sonication-sensitive cells at $T_0$ and sonication-resistant spores at 48 h PS were counted microscopically using a Neubauer chamber. Mature spores at 72 h PS were quantified by subjecting samples to heat- and sonication-treatments followed by plating on nutrient agar medium and counting colonies after 5 days at 32°C. Values indicate the average of at least three biological replicates and one standard deviation.

**Table S2 Fold change in transcript levels in a *fruA* mutant relative to a WT strain for genes in late-acting operons based on McLoon *et al.* 2021**

| Gene | 12 h[a] | 24 h[a] | MXAN_# | MXAN_RS# |
|---|---|---|---|---|
| *exoA* | 420 | 3.2 | 3225 | 15620 |
| *exoB* | 680 | | 3226 | 15625 |
| *exoC* | 240 | | 3227 | 15630 |
| *exoD* | 42 | | 3228 | 15635 |
| *exoE* | 11 | | 3229 | 15640 |
| *exoF* | 10 | | 3230 | 15645 |
| *exoG* | 16 | | 3231 | 15650 |
| *exoH* | 10 | | 3232 | 15655 |
| *exoI* | 15 | | 3233 | 15660 |
| *exoL* | 5.3 | | 3259 | 15785 |
| *exoM* | 5.3 | 2.6 | 3260 | 15790 |
| *exoN* | 32 | 3.5 | 3261 | 15795 |
| *exoO* | 56 | 4.3 | 3262 | 15800 |
| *exoP* | 10 | | 3263 | 15805 |
| *nfsA* | 60 | | 3371 | 16320 |
| *nfsB* | 56 | | 3372 | 16325 |
| *nfsC* | 52 | | 3373 | 16330 |
| *nfsD* | 49 | | 3374 | 16335 |
| *nfsE* | 42 | -1.7 | 3375 | 16340 |
| *nfsF* | 97 | 3.0 | 3376 | 16345 |
| *nfsG* | 110 | 2.8 | 3377 | 16350 |
| *nfsH* | 100 | 3.5 | 3378 | 16355 |
| *fadI* | 1.7 | -4.6 | 5372 | 26070 |
| *fadJ* | 2.3 | -4.0 | 5371 | 26065 |

[a]Data from Additional file 25 in (10). Positive numbers indicate fold increase in a *fruA* mutant relative to a WT strain. Negative numbers indicate fold decrease. Blank indicates no change relative to the WT strain, based on $p > 0.05$.

**Table S3  Strains, plasmids and primers**

| **Bacterial strain, plasmid or primer** | **Description** | **Source or reference** |
|---|---|---|
| **Strain** | | |
| *E. coli* | | |
| BL21(DE3) | $F^-$ *ompT hsdS*$_B$($r_B^-$ $m_B^-$) *gal dcm* with DE3, a λ prophage carrying the T7 RNA polymerase gene | Novagen |
| DH5α | $\lambda^-$ ϕ80dlacZΔM15 Δ(*lacZYA-argF*)*U169 recA1 endA1 hsdR17*($r_K^-$ $m_K^-$) *supE44 thi-1 gyrA relA1* | (11) |
| *M. xanthus* | | |
| AG306 | *nla6*::pNBC6 ($Km^r$) | (12) |
| AG1152 | *exoL*::pKG52 ($Km^r$) | (6) |
| DK1622 | Laboratory strain | (13) |
| DK5208 | *csgA*::Tn*5-132* Ω 205 ($Tc^r$) | (14) |
| DK5285 | *fruA*::Tn*5 lac* Ω4491 ($Km^r$) | (15) |
| DK10524 | *exoC*::Tn*5 lac* Ω7536 ($Km^r$) | (16) |
| DK11209 | Δ*devS* | (17) |
| LS3950 | *fadI*::plasmid insertion ($Km^r$) | (18) |
| MRR7 | Δ*devI* | (19) |
| MSS2 | Δ*mrpC* MXAN_0018-MXAN_0019::pSS10 ($Tc^r$) | This study |
| MSS3 | *csgA*::pRR028 ($Km^r$) MXAN_0018-MXAN_0019:: pSS10 ($Tc^r$) | (7) |
| MSS5 | *csgA*::pRR028 ($Km^r$) MXAN_0018-MXAN_0019::pSS9 ($Tc^r$) | (7) |
| MSS10 | *nla6*::pSS11 ($Tc^r$) | This study |
| SW2808 | Δ*mrpC* | (20) |
| PH1200 | *Δ*(*nfsA-H*) | (21) |
| **Plasmid** | | |
| pET11a/FruA-DBD-$H_8$ | $Ap^r$; pET11a with a gene encoding FruA-DBD-$His_8$ under control of a T7 RNA polymerase promoter | (22) |
| pET11km/FruA-$H_6$ | $Km^r$; pET11km with a gene encoding FruA-$His_6$ under control of a T7 RNA polymerase promoter | S. Inouye; (23) |
| pMAL-c5x/MBP-ΔCNla6 DBD | $Ap^r$; pMAL-c5x with a fragment of *nla6* designed to encode the Nla6 DBD, but lacking a CG bp | M. Ma and A. Garza |
| pMAL-c5x/MBP-Nla6 DBD | $Ap^r$; pMAL-c5x with a gene encoding MBP-Nla6 DBD under control of the *tac* promoter | This study |
| pMR3487 | $Tc^r$; *M. xanthus* 1.38-kb-$P_{IPTG}$- MCS_A-PR4::*lacI* | (24) |
| pMR3691 | $Tc^r$; *M. xanthus* MXAN_0018-MXAN_0019-$P_{R3-4}$::*vanR*-$P_{van}$-MCS_G | (24) |
| pPH158 | $Km^r$; pET28a *mrpC* (pET28a with a gene encoding $His_6$-MrpC under control of a T7 RNA polymerase promoter) | (25) |
| pPV391 | pCR2.1-TOPO with *dev* DNA spanning positions -321 to +71 | (26) |
| pSS9 | $Tc^r$; pMR3691 with *fruA D59E* inserted at MCS_G | (7) |
| pSS10 | $Tc^r$; pMR3691 with *fruA* inserted at MCS_G | (7) |
| pSS11 | $Tc^r$; pMR3487 with a fragment of *nla6* (+81 to +699 relative to the translation start codon); used to create a plasmid | This study |

| | | |
|---|---|---|
| | insertion mutation in *M. xanthus.* DK1622 DNA using primers Nla6 Fwd and Nla6 Rev. The fragment was joined to pMR3487 using Gibson Assembly (27). | |
| pSS14 | $Tc^r$; pMR3487 with a fragment including the *nfsA* upstream region (-290 to +83 relative to the translation start codon); used to | This study |
| **Primer** | **Description** | **Source or reference** |
| 16S rRNA fwd | CAAGGGAACTGAGAGACAGG | (28) |
| 16S rRNA rev | CTCTAGAGATCCACTACTTGCG | (28) |
| 3487 seq Fwd1 | GTAAAAAGGCCGCGTTGCTGG | This study |
| 3487 seq Fwd2 | CCTTTGATCTTTTCTACGGGG | This study |
| 3487 seq Fwd3 | GTCCATTCCGACAGCATCGCC | This study |
| 3487 seq Fwd4 | ACCAAACGTTTCGGCGAGAAG | This study |
| 3487 seq Fwd5 | CTGGATACCGCGCGGCTCAAG | This study |
| *exoA-I* -171 fwd | CCAGCCCCGGGAAATGGGAAG | This study |
| *exoA-I* -1 rev | CCTTGGATCGCAGTGGGTTAC | This study |
| *exoL-P* -217 fwd | CAAGATGGTGGCCTGGATG | This study |
| *exoL-P* +56 rev | CCTTGCCCGTCGCCATTCACG | This study |
| exoA-NF4 | CAGCAAGGGCGGACAGAT | (29) |
| exoA-NR4 | CGGAGCATGACCTCGTGT | (29) |
| *fadIJ* -214 fwd | CCGCGAAGTTCCTGGTGGAGG | This study |
| *fadIJ* -5 rev | CATGTGTGCCACACCTCCAGC | This study |
| LK1298 | CGAGGACCAGCGCTCGTC | (26) |
| LK1331 | CCAAGCTTGCTCACGTTGCAGACGGGG | (26) |
| mbp fwd colpcr | GTCGATGAAGCCCTGAAAG | This study |
| MBP-Nla6 add C fwd | CTCGGGATCGAGGGAAGGCACACCTTCCGGCTGCCCTTC | This study |
| MBP-Nla6 add C rev | AGGGCAGCCGGAAGGTGTGCCTTCCCTCGATCCCGAG | This study |
| Mxan_5372 F1 | CTGGAGTCTTCACGGACGAT | (29) |
| Mxan_5372 R1 | TCTGTTCGACAACGAGGTCA | (29) |
| Mxan_3259 F3 | TCCTCTCCGGGCAGAAGAC | (29) |
| Mxan_3259 R3 | GCATCGATGATCTCCGTCA | (29) |
| Nfs -290G | ATTGATTCCATTTTTACACTGATGAGGTACCGAATTCCGCTTCCGGGCCCGATTCCTC | This study |
| Nfs +83G | TCTCCTTACGCATCTGTGCGGTATTCTCGAGCCCGGGGACGGCCAACGAAGCAAAGACG | This study |
| *nfsA-H* -166 fwd | CTGCCCCGCGTGACGACC | This study |
| *nfsA-H* +35 rev | GGAGTCCGCGTCACCCGAC | This study |
| nfsA-NF | TTCTTCATCCTGGACAAGCAC | (29) |
| nfsA-NR | TCCAGGTTGACGCGGTAG | (29) |
| Nla6 DBD For | CACACCTTCCGGCTGCCCTTCG | M. Ma and A. Garza |
| Nla6 DBD Rev | ACCTGCAGGTCAGCGGATGACGAAC | M. Ma and A. Garza |
| Nla6 Fwd | ATTGATTCCATTTTTACACTGATGAGGTACCGAATTCTGACACAAGGTCGAGATCGCATT | This study |
| Nla6 Fwd 4 | GGGCATGCGCAAGGTCATCGA | This study |

| | | |
|---|---|---|
| Nla6 Fwd 5 | GAGATCGCATTCGACGGGCATG | This study |
| Nla6 Rev | TCTCCTTACGCATCTGTGCGGTATTCTCGAGCCCGGGTCACATCTCGAACACGCCGGG | This study |
| PMR3487G Fwd | AATACCGCACAGATGCGTAA | This study |
| PMR3487G Rev | TCATCAGTGTAAAAATGGAATCAATAAA | This study |
| PMR3487 Rev | CCTTTTGCTGGCCTTTTGCTCACA | This study |
| pMR3691 MCS G-F | CACGATGCGAGGAAACGCA | (7) |
| pMR3691 MCS G-R | CACCGGTACGCGTAACGTTC | (7) |

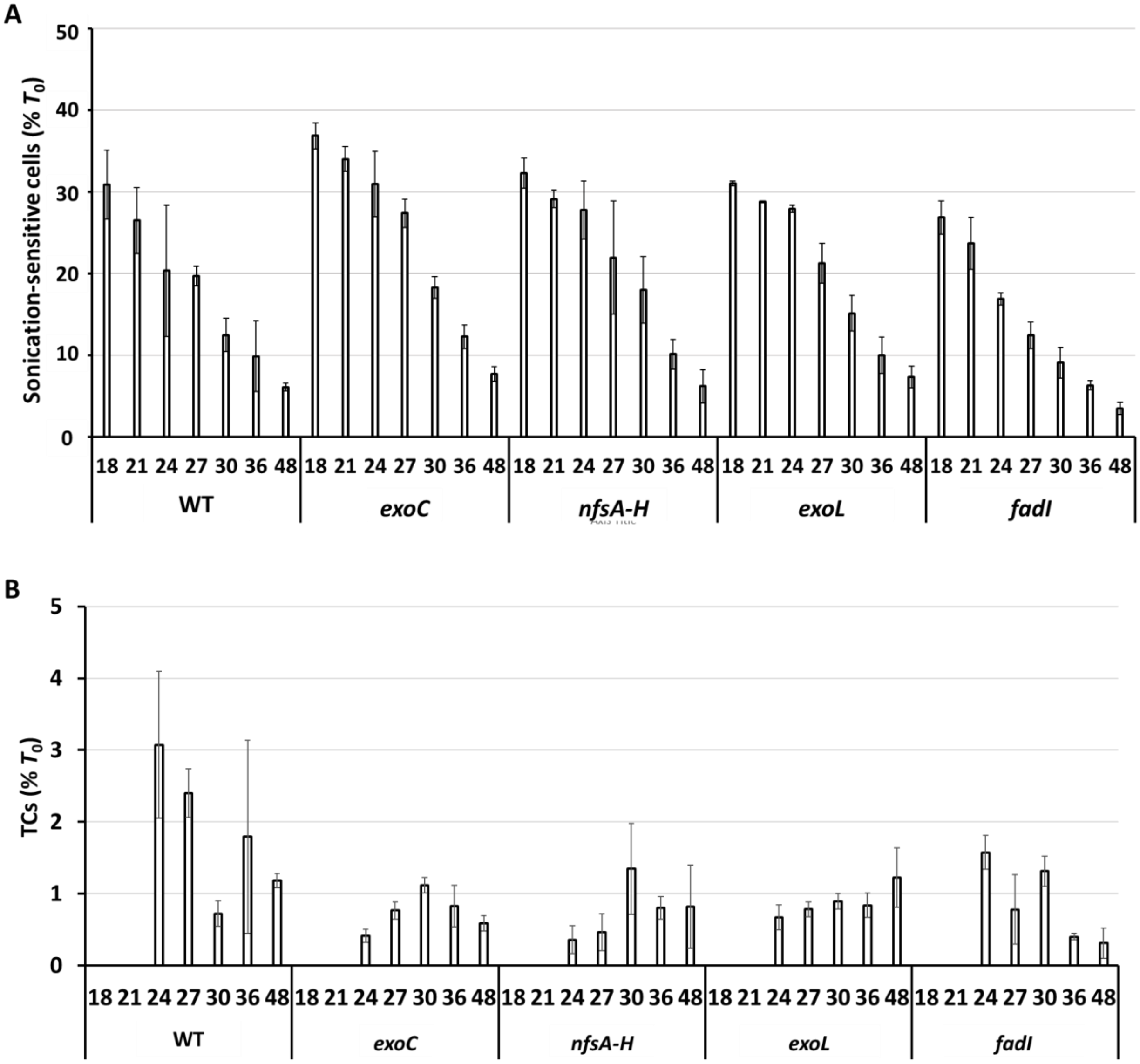


Figure S1. Sonication-sensitive cells during *M. xanthus* development. The WT strain and its mutant derivatives were subjected to starvation under submerged culture conditions. Samples were collected at the indicated times PS for quantification of total sonication-sensitive cells (A) and TCs (B). Values are expressed as a percentage of the number of rod-shaped cells present at the time starvation initiated development ($T_0$) (Table S1). Bars show the average of three biological replicates and error bars indicate one standard deviation.

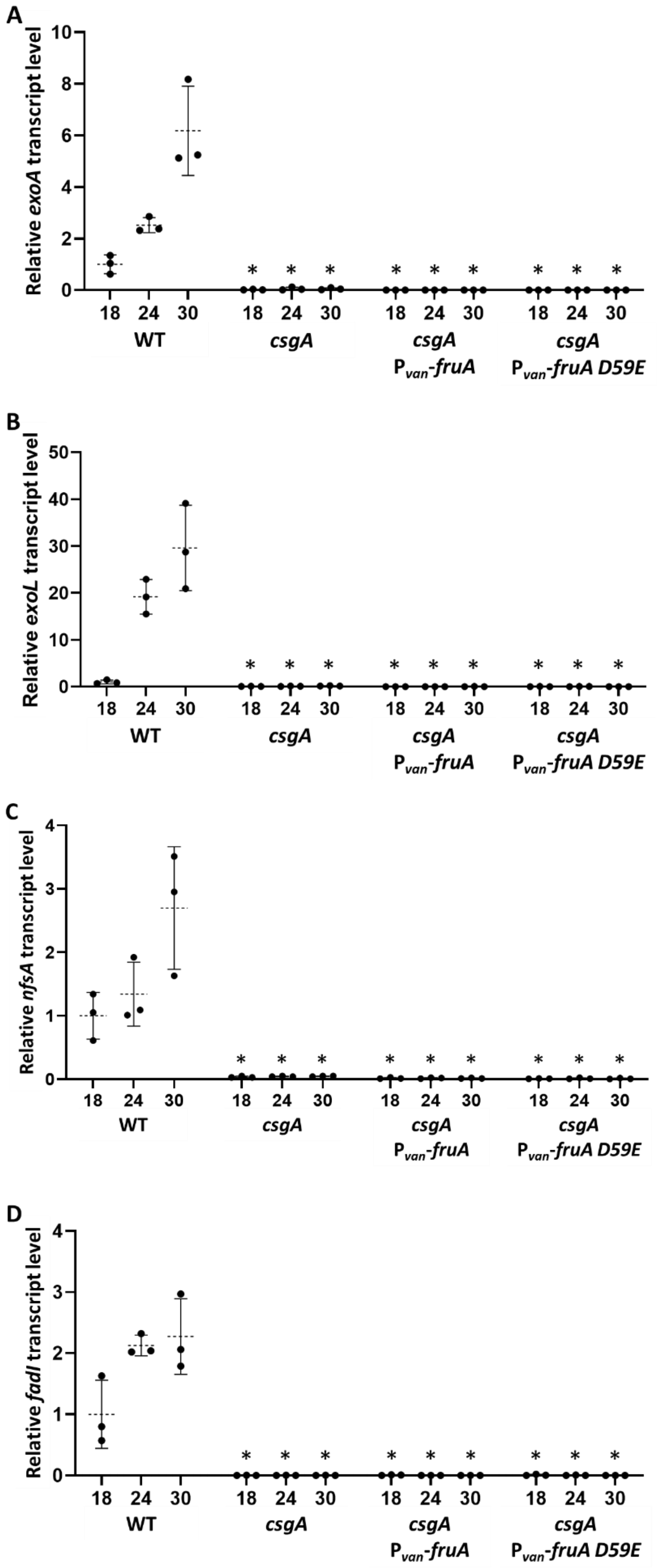
A
Relative exoA transcript level
0
2
4
6
8
10
B
Relative exoL transcript level
0
10
20
30
40
50
C
Relative nfsA transcript level
0
1
2
3
4
D
Relative fadI transcript level
0
1
2
3
4
*
18
24
30
WT
csgA
csgA
P$_{van}$-fruA
csgA
P$_{van}$-fruA D59E

Figure S2. Transcript levels in the WT strain, the *csgA* mutant, and the *csgA* mutant with inducible *fruA* or *fruA D59E* late in *M. xanthus* development. The WT strain and its mutant derivatives were subjected to starvation under submerged culture conditions. Samples were collected at the indicated times PS for measurement of the *exoA* (A), *exoL* (B), *nfsA* (C), and *fadI* (D) transcript levels by RT-qPCR. Expression of *fruA* or *fruA D59E* fused to $P_{van}$ was induced with vanillate (0.5 mM) during growth and at 0 h PS. Graphs show the data points and the average of three biological replicates, relative to the WT strain at 18 h PS, and error bars indicate one standard deviation. Asterisks indicate a difference ($p < 0.05$ in Student's two-tailed *t*-tests) from the WT strain at the corresponding time PS.

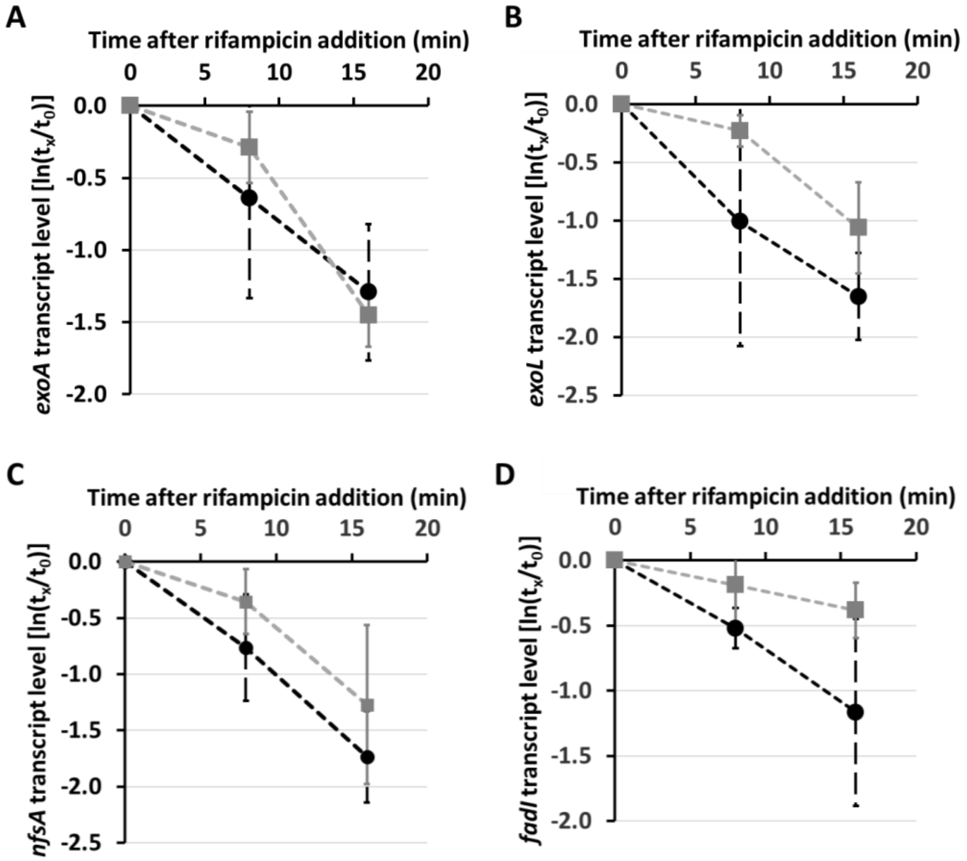


**E**

| Transcript | Half-life in the WT strain (min) | Half-life in the *mrpC* mutant (min) | *p* value |
|---|---|---|---|
| *exoA* | 6 ± 4 | 11 ± 5 | $p = 0.35$ |
| *exoL* | 8 ± 2 | 12 ± 5 | $p = 0.20$ |
| *nfsA* | 7 ± 0.4 | 7 ± 1 | $p = 0.21$ |
| *fadI* | 8 ± 2 | 12 ± 5 | $p = 0.20$ |

Figure S3. Transcript stability in the WT strain and the *mrpC* mutant midway in *M. xanthus* development. The WT strain and the *mrpC* mutant were subjected to starvation under submerged culture conditions for 18 h. The overlay was replaced with fresh starvation buffer containing rifampicin (50 μg/mL) and samples were collected immediately ($t_0$) and at the indicated times ($t_x$) for measurement of the *exoA* (A), *exoL* (B), *nfsA* (C), and *fadI* (D) transcript levels by RT-qPCR. Transcript levels at $t_x$ were normalized to that at $t_0$ for each of three biological replicates and used to determine the transcript half-life for each replicate. The graph shows the average $\ln(t_x/t_0)$ and one standard deviation for the three biological replicates of the WT strain (black dashed line) and the *mrpC* mutant (gray solid line). The average half-life and one standard deviation, as well as the *p* value from a Student's two-tailed *t*-test, are reported in (E).

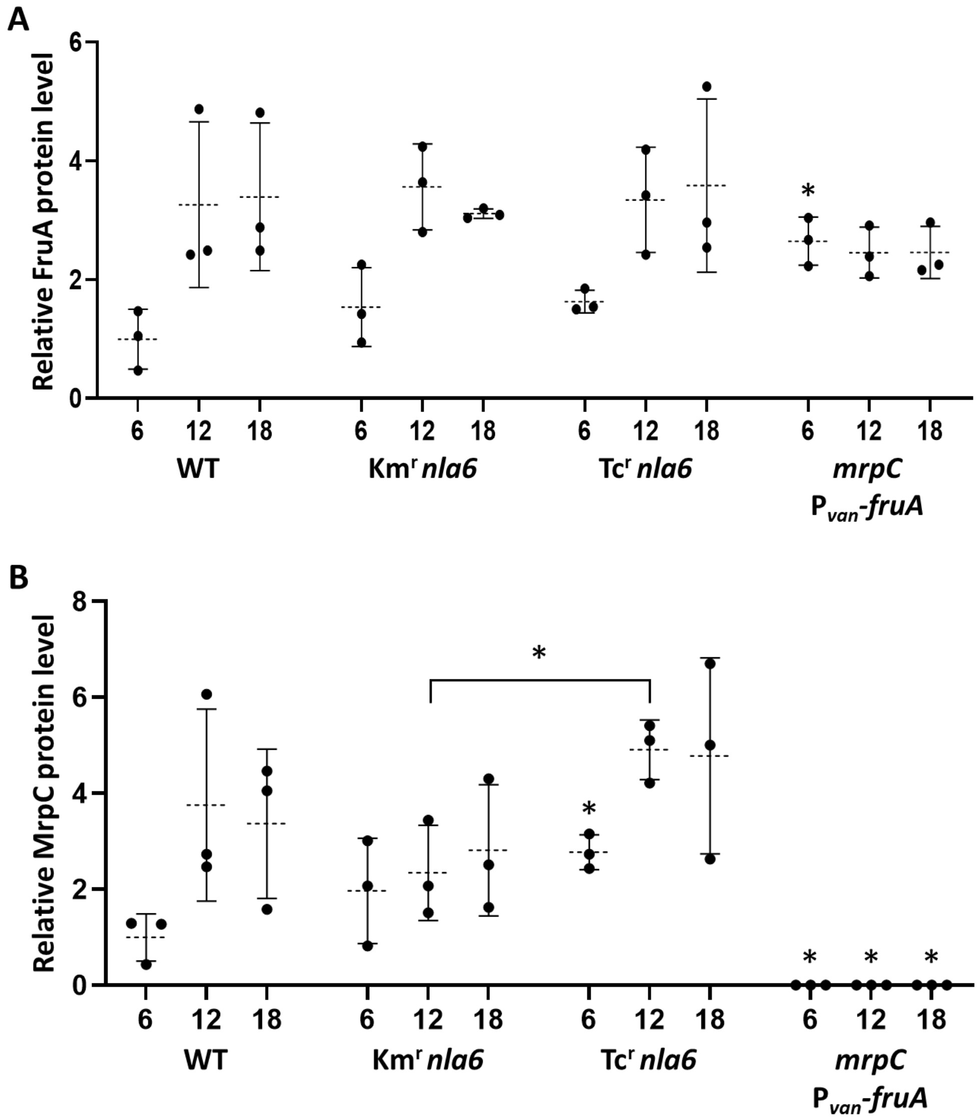


Figure S4. FruA and MrpC levels early in *M. xanthus* development. The WT strain and its indicated mutant derivatives were subjected to starvation under submerged culture conditions. Samples were collected at the indicated times PS for measurement of FruA (A) and MrpC (B) levels by immunoblot analysis. Expression of *fruA* fused to $P_{van}$ was induced with vanillate (0.5 mM) during growth and at 0 h PS. Graphs show the data points and the average of three biological replicates, relative to the WT strain at 6 h PS, and error bars indicate one standard deviation. Asterisks above error bars indicate a difference ($p < 0.05$ in a Student's two-tailed *t*-test) from the WT strain at the corresponding time PS and the asterisk above the bracket indicates a difference between the mutants.

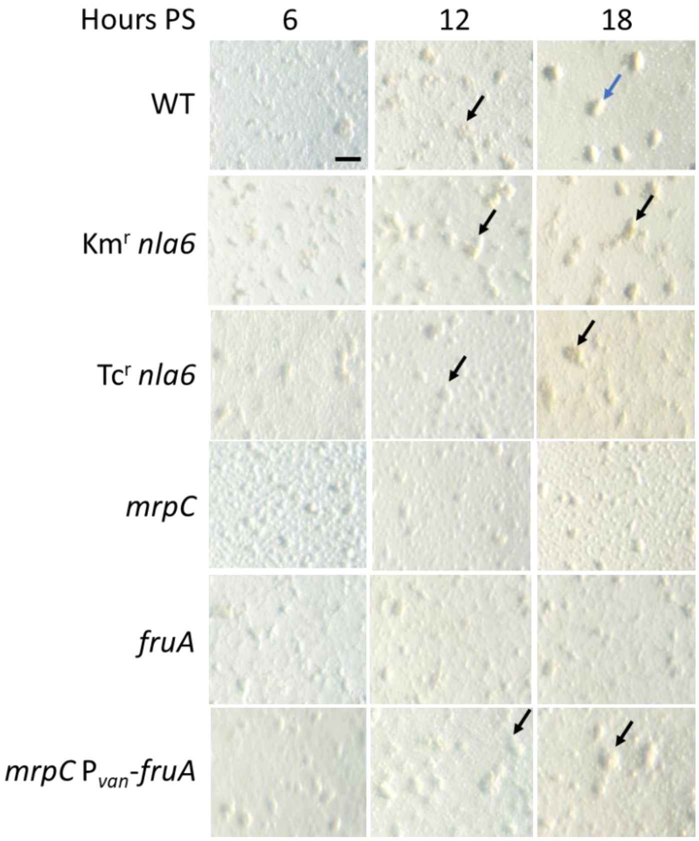


Figure S5. Early development of *M. xanthus* strains. The WT strain and its indicated mutant derivatives were subjected to starvation under submerged culture conditions and microscopic images were obtained at the indicated times PS. Expression of *fruA* fused to $P_{van}$ was induced with vanillate (0.5 mM) during growth and at 0 h PS. The WT strain, both *nla6* mutants, and the *mrpC* $P_{van}$-*fruA* mutant formed nascent mounds by 12 h (black arrows); however, only the WT strain formed compact mounds by 18 h (blue arrow). The *mrpC* and *fruA* mutants failed to form mounds. Bar, 100 μm. Similar results were observed in at least three biological replicates.

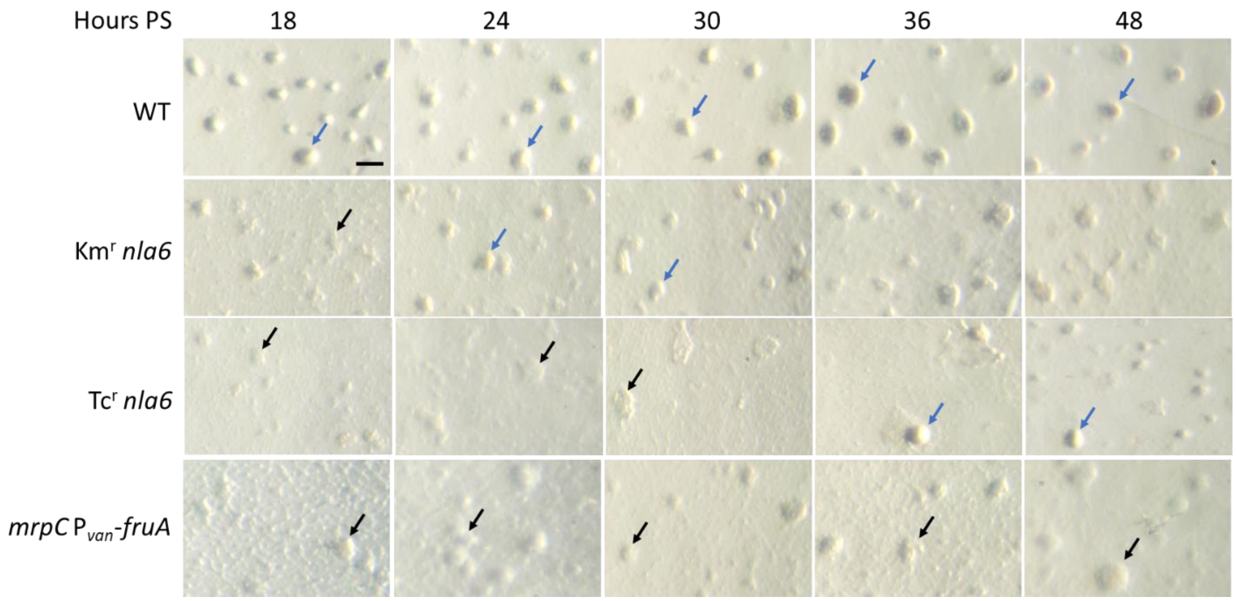


Figure S6. Late development of *M. xanthus* strains. The WT strain and its indicated mutant derivatives were subjected to starvation under submerged culture condition and microscopic images were obtained at the indicated times PS. Expression of *fruA* fused to $P_{van}$ was induced with vanillate (0.5 mM) during growth and at 0 h PS. The WT strain formed compact mounds (blue arrows) by 18 h, which darkened by 36 h. The $Km^r$ *nla6* mutant formed nascent mounds (black arrows) by 18 h and compact mounds at 24 and 30 h, but the mounds failed to darken and became less compact at 36 and 48 h. The $Tc^r$ *nla6* mutant formed nascent mounds by 18 h, but the mounds did not become compact until 36 h and did not darken by 48 h. The *mrpC* $P_{van}$-*fruA* mutant formed nascent mounds by 18 h, but the mounds did not become compact and did not darken by 48 h. Bar, 100 μm. Similar results were observed in at least three biological replicates.

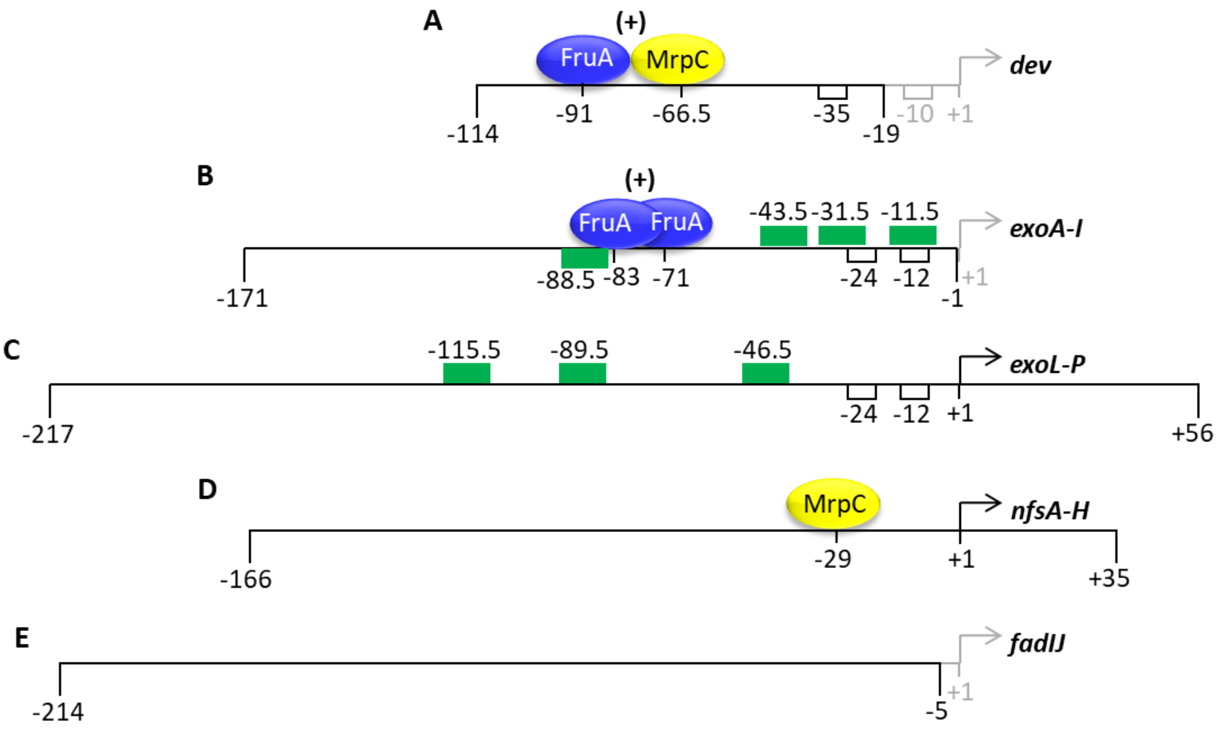


Figure S7. Features of promoter region DNA fragments used in DNA-binding assays. (A) The *dev* promoter region fragment spans from -114 to -19 relative to the transcriptional start site (bent arrow at +1) (17) and is bound cooperatively by FruA and MrpC to sites centered at -91 and -66.5, respectively, which regulate transcription positively (+) (26, 30). (B) The *exoA-I* fragment (-171 to -1) was shown to be bound by at least three FruA DBDs, and DNase I footprinting revealed a protected region spanning at least from -89 to -64, which may include two binding sites centered at -83 and -71 that overlap by 3 bp and regulate transcription positively (22). The fragment is also bound by MBP-Nla6 DBD and contains four DNA sequences (green rectangles centered at the indicated positions) similar to the half-site consensus sequence for Nla6 binding (6). (C) The *exoL-P* fragment (-217 to +56) is also bound by MBP-Nla6 DBD and contains three DNA sequences similar to the Nla6 half-site consensus sequence (6). (D) The *nfsA-H* fragment (-166 to +35) was shown to be bound by MrpC at a site predicted to be centered at -29 (5). (E) The *fadIJ* fragment (-214 to -5) was not known to be bound directly by FruA or MrpC, but mutations in *fruA* or *mrpC* prevent developmental expression of a reporter transcriptionally fused to a *fadIJ* upstream DNA fragment (-340 to -4, based on the transcriptional start site depicted) (18). The transcriptional start sites depicted for *fadIJ*, *nfsA-H*, and *exoL-P* were determined by Cappable-seq (31) (note: the site with maximum counts at 24 h PS was chosen), while those depicted for *dev* (17) and *exoA-I* (designated $P_{D1}$) (22) were determined by primer extension analysis at high and low resolution, respectively, and were at +2 and -51 relative to the Cappable-seq sites with maximum counts at 24 h PS (31).

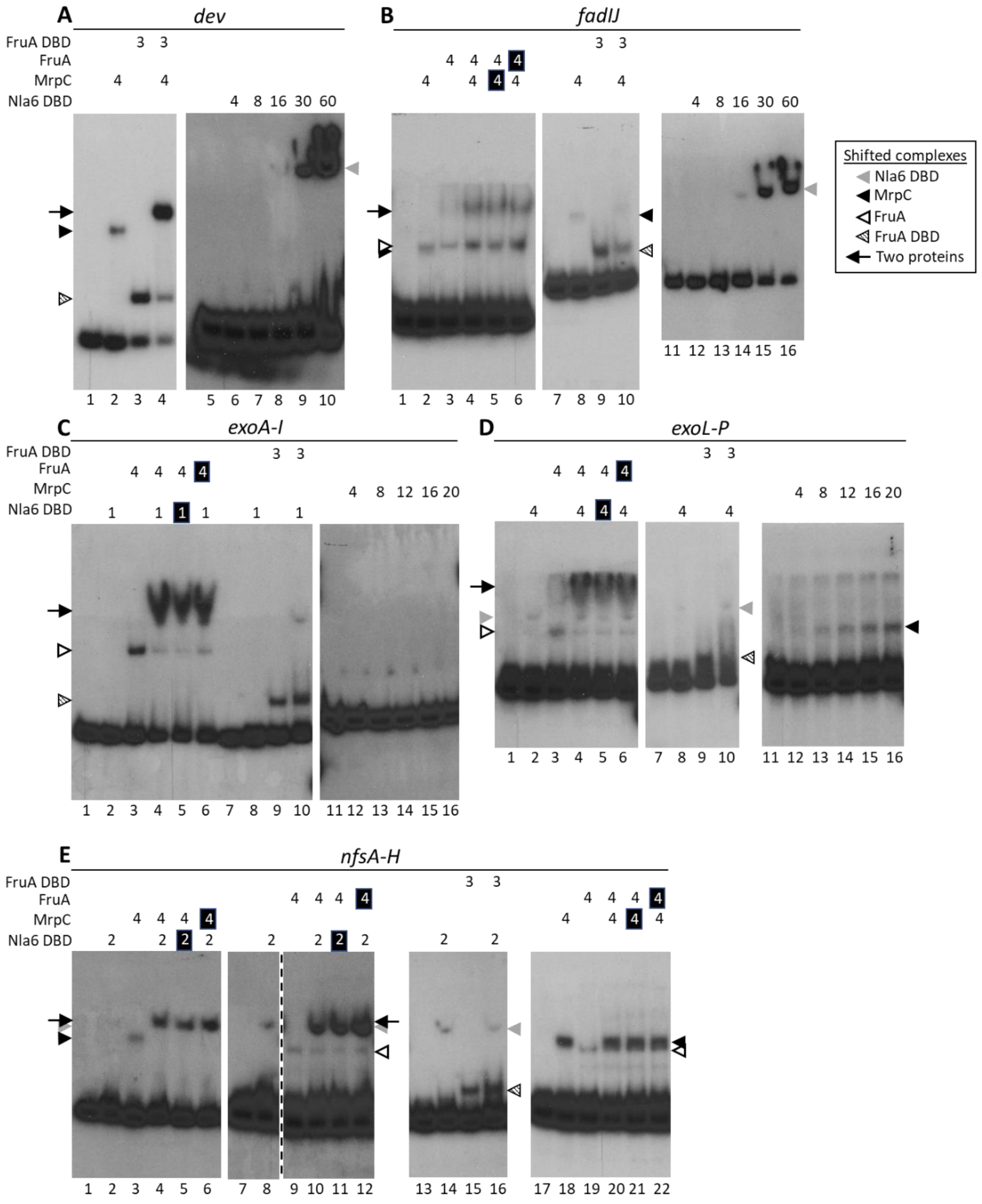


Figure S8. Binding of the Nla6 DBD, MrpC, FruA, and the FruA DBD to the promoter regions of the late-acting operons. EMSAs were performed with $^{32}$P-labeled DNA fragments (2 nM) of the *dev* (-114 to -19) (A), *fadIJ* (-214 to -5) (B), *exoA-I* (-171 to -1) (C), *exoL-P* (-217 to +56) (D), and *nfsA-H* (-166 to +35) (E) promoter regions, and MBP-Nla6 DBD, $His_6$-MrpC, FruA-$His_6$, and FruA DBD-$His_8$ at the $\mu$M concentrations indicated by numbers above each autoradiographic image.

A white number on a black background indicates a protein that was added to the DNA-binding reaction 10 min later than the other protein. The positions of migration of shifted complexes are indicated along the side of each image by arrowheads for binding of individual proteins and by arrows for binding of two proteins (see boxed key). A dashed line indicates that intervening lanes were cropped from the autoradiographic image. Lanes are numbered below images.

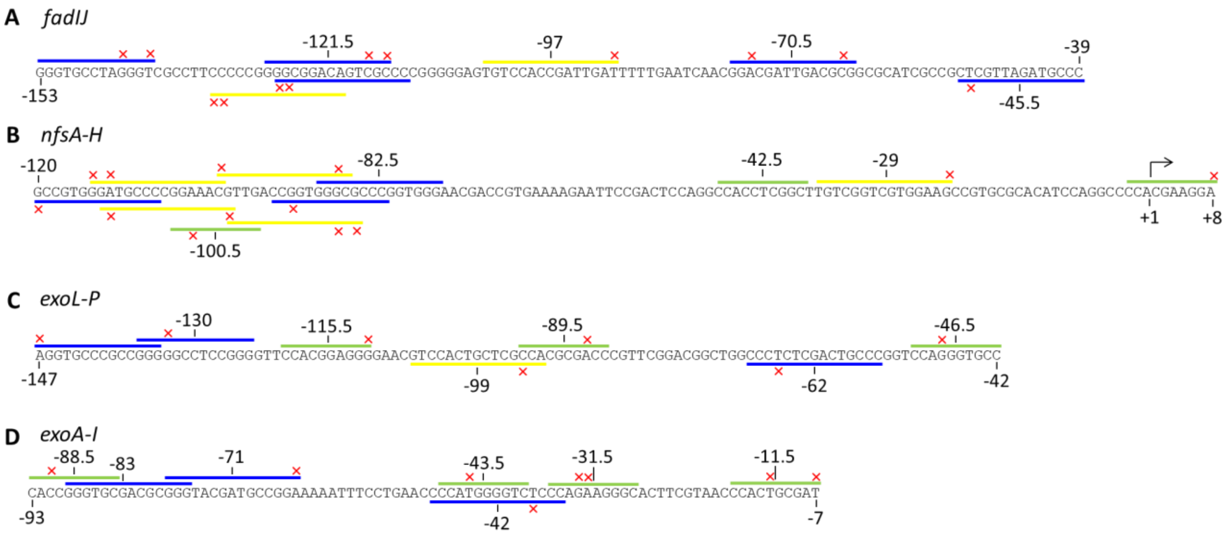


Figure S9. DNA sequences in the promoter regions of late-acting operons match consensus sequences for binding of MrpC, FruA, and Nla6. DNA sequences matching the MrpC (TGTYN$_8$RAC) (5), FruA (GGGYRN$_{4-6}$YGGG) (30), and Nla6 half-site [C(C/A)ACGN$_2$GNC] (6) consensus sequences are indicated by yellow, blue, and green lines, respectively, above (strand shown) or below (strand not shown) the promoter regions. A maximum of two mismatches to consensus sequences were allowed and are indicated by a red x. Numbers indicate positions relative to transcriptional start sites shown in Figure S7.
